# MYC-induced oncogenesis is dependent on acidic patches within its N-terminal intrinsically disordered domain

**DOI:** 10.1101/2025.06.16.659507

**Authors:** Victor Llombart, David O’Connor, Jonas Demeulemeester, Amandeep Bhamra, Silvia Surinova, Adam Turna, Kent Fung, Lingyi Wang, Yang Li, Tanya Rapoz-D’Silva, Farah Ahmed, Henri Niskanen, Ida Stöppelkamp, Denes Hnisz, Sandro Bottaro, Carlo Fisicaro, Shuning He, A Thomas Look, Marc R Mansour

**Affiliations:** Department of Haematology, UCL Cancer Institute, University College London, London, UK; Proteomics Research Translational Technology Platform, UCL Cancer Institute, University College London, London, UK; Developmental Biology and Cancer, UCL Great Ormond Street Institute of Child Health, London, UK; The Francis Crick Institute, London, UK; Department of Oncology, KU Leuven, Leuven, Belgium; VIB Center for Cancer Biology, Leuven, Belgium; Max Planck Institute for Molecular Genetics, Berlin, Germany; Peptone Ltd, Corporate Headquarters, London, UK; Department of Pediatric Oncology, Dana-Farber Cancer Institute, Harvard Medical School, Boston, MA; Division of Pediatric Hematology/Oncology, Boston Children’s Hospital, Boston, MA, USA

## Abstract

MYC is one of the most enticing therapeutic targets for cancer but clinical-grade inhibitors are still lacking. By site-saturation mutagenesis screening, we identified several evolutionarily conserved acidic patches within the intrinsically-disordered MYC N-terminus that were confirmed to be functionally essential in different cell models and in vivo. Beyond modulating MYC’s global transcriptional activity, these negatively charged patches regulate the interaction with chromatin-modifying complexes including those with histone acetyl-transferase activity. One of the key interactions is established with the co-factor TRRAP, a subunit shared between several Histone Acetyl-Transferase complexes. The protein-protein binding between MYC and TRRAP predominantly relies on two of the N-terminal negative clusters that are located outside MYC-Box-II (MBII) and drive oncogenesis. Our work identifies a new multivalent MYC subdomain that presents new therapeutic vulnerabilities providing invaluable insights for the development of new therapeutic approaches.

## INTRODUCTION

MYC is a pleiotropic transcription factor that regulates core cellular processes such as growth, metabolism, stemness, apoptosis and cell differentiation (1–4). In haematopoiesis, MYC also plays a major role by regulating self-renewal and differentiation of haematopoietic stem cells (5). Multiple lines of evidence link aberrant MYC expression with the “hallmark” features of cancer (6, 7), with a role as a driver of both tumour initiation and maintenance across multiple different tumour subtypes (8–10).

T-cell acute lymphoblastic leukaemia (T-ALL) is a prototypic MYC-driven malignancy, activated primarily through aberrant NOTCH signalling that occurs as a result of activating *NOTCH1* mutations and/or duplication of the NOTCH-MYC-enhancer (11–14). T-ALL results from clonal expansion and uncontrolled proliferation of malignant T-cell precursors. Despite significant advancements in treatment modalities, there is still an urgent need to explore novel therapeutic strategies, particularly in relapsed or refractory cases in which the prognosis remains poor (15). Preclinical studies conducted in a variety of in vivo cancer models, including T-ALL, have shown that inhibition of MYC leads to tumour regression through differentiation and apoptosis, while anti-proliferative effects on normal tissues remain reversible (16–18). This suggests that targeting MYC has the potential to exploit a clinically meaningful therapeutic window. Unfortunately, the development of clinical grade specific MYC inhibitors still poses an immense challenge and only the synthetic peptide Omomyc and the molecular glue WBC100 have thus far reached the clinical trial stage (NCT04808362 and NCT05100251, respectively).

Understanding the structure of the MYC protein is essential for unravelling its functional complexity and designing targeted therapies. MYC consists of several functional protein domains. The C-terminal domain comprises a basic helix-loop-helix/leucine zipper (bHLH/LZ) involved in MYC:MAX heterodimerization and in DNA binding. The N-terminal transactivation domain is known to modulate the interaction with a myriad of co-factors that include transcriptional activators, repressors and chromatin modifiers that influence MYC’s activity and stability (19–22). While this regulatory domain is integral to MYC’s oncogenic function, it is intrinsically disordered (ID), making in silico design of inhibitory small molecules extremely difficult (23). However, studies on c-MYC and its paralog N-MYC, show that subregions of the N-terminus can adopt an ordered conformation when co-crystallised with a direct binding partner such as WDR5 (24) or Aurora-A (25). This is consistent with the ‘coupled folding and binding’ model, whereby many ID proteins become structured upon binding to a partner, with the energy from specific interactions compensating for the entropic penalty from ordering (26).

To identify new potential therapeutic vulnerabilities on MYC, we have generated a single amino acid resolution functional map of the MYC N-terminus using T-ALL as a prototypic MYC-dependent cancer model and identified a series of acidic amino acid residues that are grouped in patches along the MYC N-terminus. Notably, mutagenesis of these acidic clusters completely abrogates MYC oncogenic function in different cancer cell models and in vivo. Here, we investigate the functional implications of this diffuse subdomain and, by combining protein proximity labelling with mass spectrometry, identify critical MYC interacting partners that are dependent on acidic patches outside the classic MYC box (MB) homology regions. Understanding the interplay between MYC and its cofactors has the potential to open promising avenues for the development of novel anti-MYC therapies in T-ALL and other malignancies. Our findings also highlight the biological relevance of acidic domains on oncogenicity, with implications for the myriad of other TFs containing such domains.

## RESULTS

### Site saturation mutagenesis screening identifies N-terminal acidic residues as critical for MYC function

The protein sequence of MYC contains several evolutionarily-conserved segments that modulate MYC function including the MYC homology boxes (MBs) and the basic helix-loop-helix leucine-zipper (bHLH-LZ) C-terminal domain (Figure 1A). To perform structure-function experiments, we utilised a c-MYC-dependent T-ALL cell line established from mice where, on the addition of doxycycline, expression of endogenous MYC is repressed leading to cell death (4188 cells, Figure 1B; Supplemental figure 1) (27). In these cells, ectopic expression of MYC through retroviral transduction can completely rescue cell viability in the presence of doxycycline (Fig 1 C and 1D).

**Figure 1:**
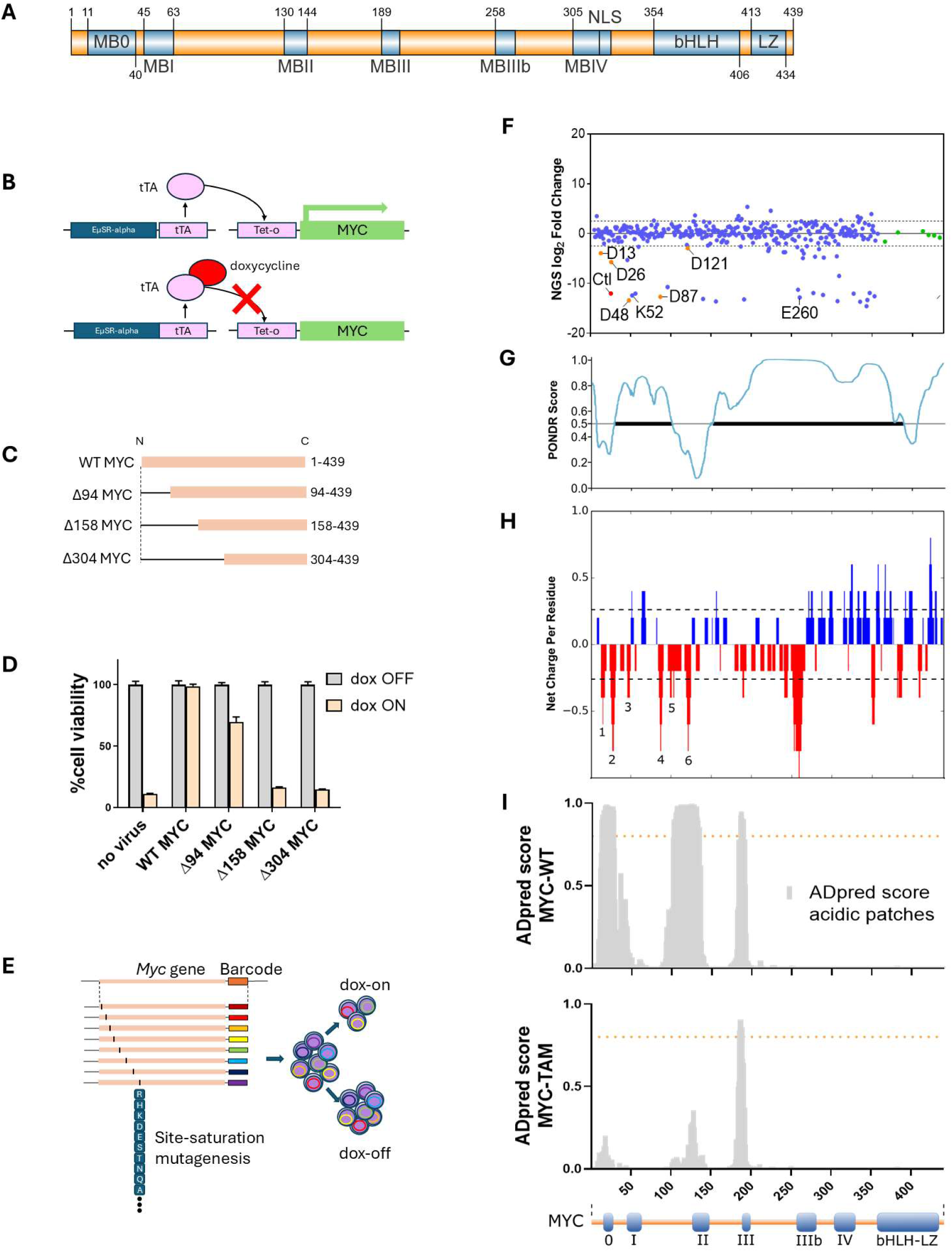
Single amino acid resolution screening of MYC in T-ALL cells. A) Schematic of the MYC protein. High homology MYC boxes (MBs) 0–IV and the basic helix-loop-helix leucine zipper (bHLH-LZ) domain are shown in blue (generated using IBS2.0 (85). B) Structure of the MYC gene in murine 4188 cells. Tetracycline trans-activating protein (tTA) is expressed under the control the immunoglobulin heavy chain enhancer and the SRa promoter (EµSR-alpha-tTA). When expressed, tTA controls the transcription of the MYC transgene placed under the control of the tetracycline-responsive promoter (Tet-o). The presence of doxycycline represses transcription mediated by tTA and results in inhibition of MYC expression. C) Schematic of the MYC deletion mutants screened. Δ94 lacks residues 1-94, Δ158 lacks residues 1-158 and Δ304 lacks 1-304. D) Cell viability of 4188 cells expressing MYC-WT (wild-type) or MYC deletion mutants and incubated 72h with doxycycline (20ng/mL). E) Schematic of the experimental workflow for generating a single-amino acid MYC mutant library and the screening of drop-out variants on 4188 cells after doxycycline incubation. Each amino acid position was barcoded to ease the library deconvolution after doxycycline treatment. F) Dot plot showing the site saturation mutagenesis screening results after 96 hrs of doxycycline treatment. Barcode reads were obtained by next-generation sequencing and log2 of the ratio dox^on^ versus dox^off^ is shown. G) Plot depicting intrinsic disorder for MYC. PONDR (Predictor of Natural Disordered Regions) VSL2 score is shown on the y axis, and amino acid positions are shown on the x axis (mid panel; PONDR score>0.5 indicates disorder). H) NCPR plot for MYC showing basic (blue) and acidic (red) amino acid clusters. Acidic patches under investigation are numbered 1: D13/D15/D17, 2: D26/E27/E28/E29, 3: E47/D48/E54, 4: E85/D86/D87/D88, 5: D98/E101/E105/D110, 6: D119/D121/D122/E123. I) Predicted activation domains across MYC wild-type (MYC-WT) using *ADpred* (37). Mutation of acidic residues to alanine results in a decrease of predicted activation domain function in MYC total acidic mutant (MYC-TAM).

We first addressed the importance of the MYC N-terminus in sustaining growth of these cells by retrovirally expressing MYC constructs harbouring different length N-terminal deletions (Figure 1C and 1D). Consistent with the notion that the N-terminal transactivation domain (TAD) of MYC is essential for its biological activity (28, 29), progressive truncation of the protein’s N-terminus resulted in a gradual loss of function with the deleterious mutant MYCΔ158 (lacking residues 1-158) showing a ≈90% reduction in cell viability (Figure 1D). We confirmed that this loss-of-function phenotype was not a result of an alteration on protein expression levels (Supplemental Figure 2A and 2B), nor an effect of a loss of interaction with MYC’s obligate partner MAX (Supplementary Figure 2C).

**Figure 2:**
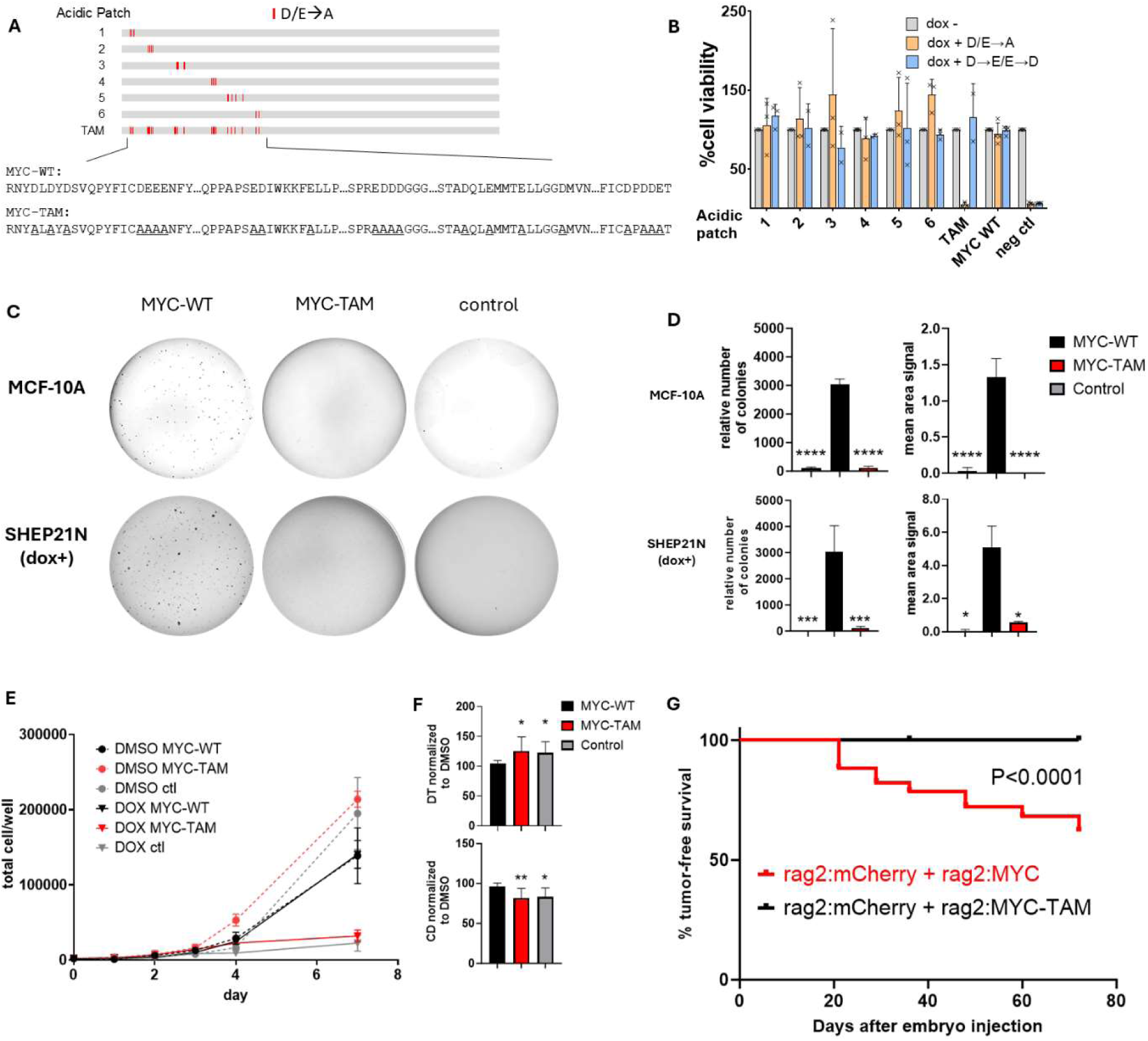
Mutation of N-terminal acidic patches abrogates MYC function in multiple cell models and impairs T-ALL initiation in vivo. A) Diagram of MYC acidic mutants tested. D or E residues were replaced by A through site-directed mutagenesis. Acidic patch 1: D13/D15/D17; Acidic patch 2: D26/E27/E28/E29; Acidic patch 3: E47/D48/E54; Acidic patch 4: E85/D86/D87/D88; Acidic patch 5: D98/E101/E105/D110; Acidic patch 6: D119/D121/D122/E123; TAM: total acidic mutant. Amino acid sequence alignment is depicted for MYC-WT (wild type) and MYC-TAM (total acidic mutant). B) Cell viability of 4188 cells expressing MYC-WT, MYC with mutations in acidic patches 1-6 (orange: charge neutralisation D/E>A and blue: charge preserving D>E or E>D) or MYC-TAM and incubated for 72h with doxycycline (20ng/mL). C) Representative images of the MCF-10A and SHEP21N (dox) cell colonies expressing MYC-WT or MYC-TAM in soft-agar plates for 2-4 weeks at 20x magnification. All colonies were stained using crystal violet. D) Bar graphs showing the number of MCF-10A relative to control (top left), SHEP21N colonies relative to control DMSO (bottom left), MCF10A colonies mean area signal (top right) and SHEP21N colonies mean area signal (bottom right). Error shown as SD. E) Cell proliferation curve of SHEP2N cells expressing ectopic MYC-WT or MYC-TAM with doxycycline (1µg/mL) or DMSO control. Non-transfected cells were used as control. F) Doubling time (DT) (top) and cell doubling (bottom) for SHEP21N cells expressing ectopic MYC-WT or MYC-TAM between day 4 and day 7. Cell doubling and doubling time are normalized to the corresponding DMSO values. Error shown as SD. G) Kaplan–Meier curves showing tumour onset in zebrafish co-injected with constructs expressing rag2:MYC-WT + rag2:mCherry (red line), versus rag2:MYC-TAM + rag2:mCherry (black line).

To identify in an unbiased fashion whether single amino acid residue vulnerabilities exist across the MYC N-terminus, we performed a site saturation mutagenesis screen where each position up to the bHLH region was mutated to all possible amino acid variants (Figure 1E; Supplementary Figure 3). To track the abundance of constructs within the library, a DNA barcode was incorporated corresponding to the position of the amino acid mutated. The library was retrovirally transduced at an MOI≤0.3 to favour incorporation of only a single MYC variant per cell. As a result, only those cells expressing a functional variant from our library would retain proliferative capacity after doxycycline-mediated inhibition of endogenous MYC. Through next generation sequencing of barcodes, we identified drop-out of the negative control (containing a premature truncation lacking the C-terminal domain) in doxycycline treated cells, as well as at residues K52 and E260, sites which have been previously functionally validated as essential for MYC function (21, 24, 30, 31) (Figure 1F). Unexpectedly, mutations of individual leucines within the LZ initially intended as negative controls were able to rescue cell growth, likely because simultaneous mutation of multiple leucines is required to fully impair function (32).

The MYC N-terminus is predicted to be highly disordered and shows a particular residue composition, rich in S, P and the acidic residues D and E, typical of intrinsically disordered regions (IDRs) (Figure 1G,; Supplementary Figure 4) (33–35). Our attention was drawn to a considerable number of acidic amino acids that scored highly in our screen, with significant drop-out of residues D13, D17, D26, D48, D87 and D121. Interestingly, acidic residues within intrinsically disordered transactivation domains have reported roles in determining the strength of activation of transcription factors, but their biological role in MYC functionality has not been tested (36). We next sought to examine the charge distribution across the whole MYC protein. The net charge per amino acid residue analysis revealed that the D and E residues within the 1-158 domain are grouped into six highly acidic patches: D13/D15/D17 (referred to hereafter as acidic patch 1), D26/E27/E28/E29 (acidic patch 2), E47/D48/E54 (acidic patch 3), E85/D86/D87/D88 (acidic patch 4), D98/E101/E105/D110 (acidic patch 5), and D119/D121/D122/E123 (acidic patch 6) (Figure 1H). The negative charge of these acidic clusters is evolutionarily conserved amongst different species from human to zebrafish, supporting an important functional role that may be relevant for MYC oncogenic activity (Supplemental Figure 5). Further supporting their potential functional relevance, some of the identified acidic patches are located within activation domains predicted by the deep learning model *ADpred* (37). Moreover, the predicted activation activity is dramatically reduced upon in-silico charge neutralization of these patches (Figure 1I).

### Acidic patches of the MYC N-terminus are required for T-ALL cell viability and leukaemia-initiation

To assess the functional role of the identified acidic patches, we individually neutralised the negative charge of each patch using site-directed mutagenesis, replacing D/E residues with alanine. We also generated a MYC total acidic mutant (MYC-TAM) by simultaneously mutating all the acidic patches to alanines (Figure 2A). Each MYC mutant was individually expressed in T-ALL 4188 cells by retroviral transduction and endogenous MYC expression was inhibited by the addition of doxycycline. None of the mutations of individual acidic patches affected MYC function (Figure 2B). The phenotypic discrepancy between the site-saturation mutagenesis screen and the alanine scanning assay is likely attributable to differences in MOI, with the MOI≤0.3 in the former assay providing a higher sensitivity for the detection of hypomorphic variants (Supplementary Figure 6). In contrast to single cluster mutations, the simultaneous charge neutralization of all six acidic patches in MYC-TAM construct resulted in a complete inability to rescue cell viability under doxycycline (Figure 2B). Notably, MYC oncogenic activity was not affected after adding mutations that preserve the negative charge of the acidic clusters (D>E or E>D), either in the individual mutants or in the MYC-TAM (Figure 2B). The functional role of MYC N-terminal acidic patches was not confined to T-ALL. Ectopic expression of MYC-TAM led to a significant decrease in colony formation of the human mammary epithelial cell line MCF-10A and the neuroblastoma human cell line SHEP-tet21/N as compared to MYC-WT (p<0.0001; Figure 2C and 2D) as well as a reduced cell proliferation (p<0.001; Figure 2E and 2F).

Aspartic and glutamic acid residues have a high net negative charge that contributes to the intrinsic disorder of proteins due to electrostatic repulsions (38). Consistently, we observed that the charge neutralization of all clusters is accompanied by a decrease in the protein disorder profile in MYC-TAM compared to MYC-WT (Supplementary Figure 7) and we asked whether the loss of MYC activity could result from this impact on the structural conformation of the N-terminus. Using the IDR from protein eIF4G2 (TIF4632_27-96_ from *S.Cereviseae*) (39), which is capable of making liquid like condensates, we generated the chimeric fusion protein 3xFlag-eIF4G2-MYC-TAM and investigated whether the oncogenic function of MYC-TAM could be rescued by restoring the disorder of the polypeptidic chain. Interestingly, we observed no effect on the capacity of the chimeric fusion protein to rescue cell viability of 4188 cells exposed to doxycycline (Supplementary figure 8) underscoring the significance of the negative charge of the MYC N-terminus as a key factor that modulates the protein activity rather than the disorder of the TAD.

Our data in cell lines indicates that the acidic clusters of the MYC N-terminus are required for maintaining tumour cell survival and proliferation, but their role in T-ALL initiation remained in question. The murine c-MYC oncogene initiates a highly aggressive T-ALL when expressed from the rag2 promoter in a T-ALL zebrafish model (40). To assess the implication of the acidic patches of the MYC N-terminus in T-ALL initiation *in vivo,* we co-injected single-cell embryos with *rag2:mCherry/rag2:Myc* or *rag2:mCherry/rag2:Myc-TAM* and monitored fish by fluorescent microscopy for tumour onset. Similar to previous observations (41), expression of MYC-WT after *rag2:mCherry/rag2:Myc* co-injection led to tumour initiation, with 38% of fish exhibiting T-ALL at 72 days. In contrast, tumour initiation was not observed when MYC-TAM was co-injected into single-cell embryos (0%, p<0.0001, Figure 2G), confirming the role of N-terminal acidic clusters on T-ALL initiation *in vivo*.

### MYC N-terminal acidic patches control transcriptional activity

To test whether the N-terminal acidic clusters play a role in modulating MYC transcriptional activity, RNA sequencing (RNA-seq) was performed after doxycycline incubation in retrovirally-transduced cells stably expressing exogenous MYC-WT, MYC-TAM and MYCΔ158. After comparing gene expression between cells in presence or absence of doxycycline, only 18 genes were significantly deregulated in MYC-WT expressing cells, reflecting effective transcriptional rescue by exogenous MYC (Figure 3A). In contrast, 676 genes were significantly deregulated in MYC-TAM cells. Cells expressing MYCΔ158 and non-transduced control cells showed 573 and 801 deregulated genes, respectively (adj. p<10^-50^, log2fold change ≥1.8). Previously reported MYC target genes such as *Cad, Ccnd2, Hspe1, Srm, Tfrc* and *Apex* (42) emerged as sensitive to doxycycline-mediated MYC inhibition in non-transduced 4188 cells as well as in cells expressing MYC-TAM and MYCΔ158, but not in MYC-WT transduced cells. Gene set enrichment analysis (GSEA) in MYC-TAM cells revealed a strong downregulation in the expression of genes from MYC targets v1 and v2 datasets (Figure 3B). Uniform Manifold Approximation and Projection (UMAP) analysis of differentially expressed genes (DEGs) revealed distinct clustering based on doxycycline treatment. Cells expressing ectopic MYC-WT with doxycycline aligned with the dox-off cluster, confirming rescue of gene expression by exogenous MYC. In contrast, cells expressing MYC-TAM or MYCΔ158 grouped with non-transduced cells, reflecting the inability of MYC-TAM to activate MYC transcriptional programs (Figure 3C). Unsupervised hierarchical clustering supported these findings (Figure 3D). A linear correlation (r² = 0.97) was observed in gene expression changes between MYC-TAM and non-transduced cells (Supplementary Figure 8). Genes most impacted by endogenous MYC depletion showed greater recovery upon ectopic MYC expression (Supplementary Figure 9). These findings highlight the critical role of MYC’s N-terminal acidic patches in orchestrating transcriptional landscapes and modulating global and specific MYC-target gene activity.

**Figure 3:**
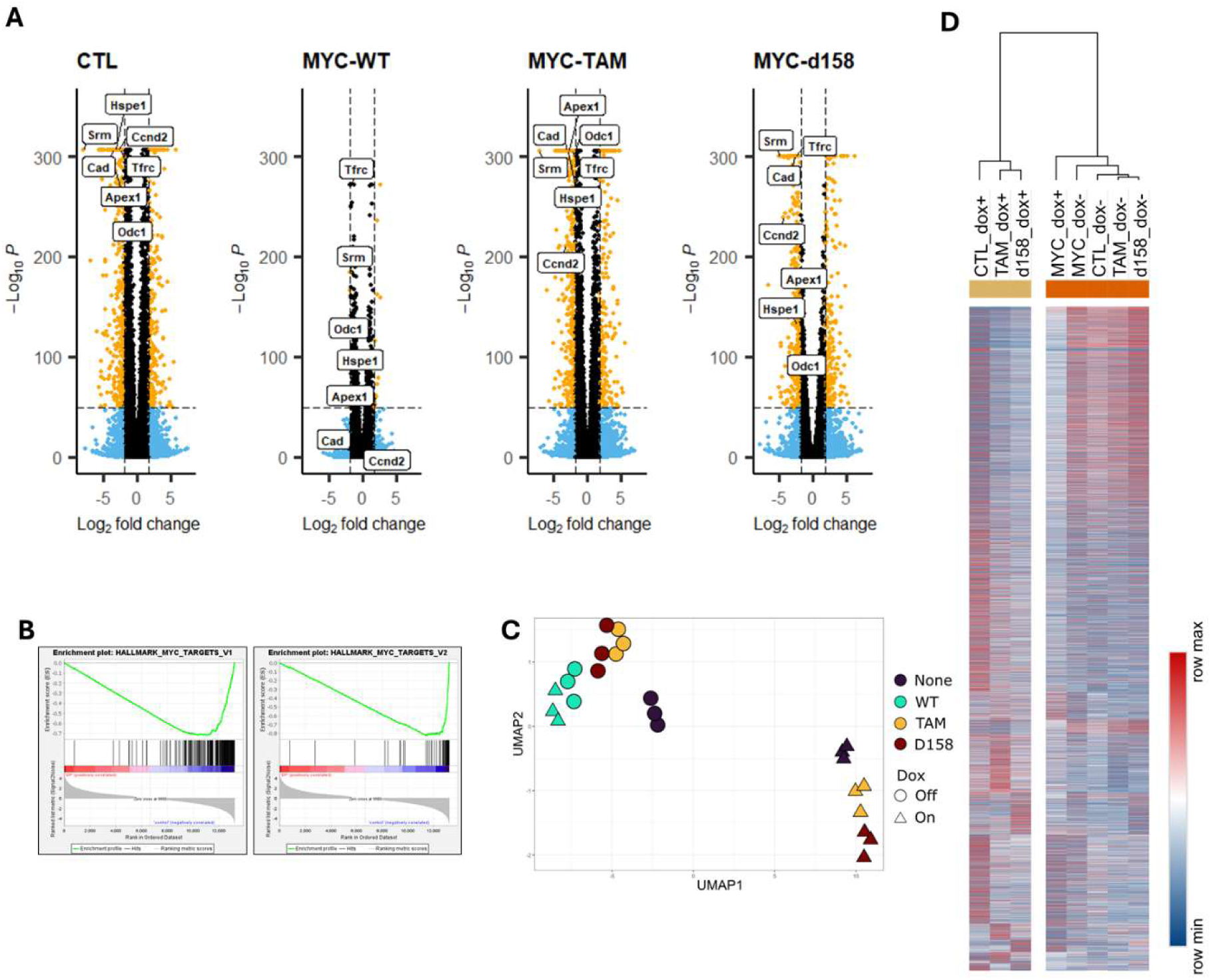
MYC N-terminal acidic patches control MYC’s global transcriptional activity. A) Volcano plots showing the differently expressed genes (DEGs) (logFC ≥ 1.8, adj. p<10e^-50^) in 4188 cells stably expressing MYC-WT, MYC-TAM, MYC-Δ158 or control (CTL, non-transduced) following doxycycline incubation. B) GSEA analysis for genes downregulated in 4188 cells stably expressing MYC-TAM incubated with doxycycline relative to no-doxycycline condition. NES, normalized enrichment score; FDR qvalue, false discovery rate q value. C) UMAP diagram showing the clustering of cell lines expressing MYC-WT, MYC-TAM or control according to DEGs in absence/presence of doxycycline. D) mRNA expression heatmap of genes from RNA-seq data of 4188 cells stably expressing MYC-WT, MYC-TAM, MYC-Δ158 or control (non-transduced) incubated with versus without doxycycline (n=3).

### The N-terminal acidic clusters do not regulate MYC protein expression, MYC:MAX interaction, protein stability, subcellular localization or DNA binding

We sought to further characterize the specific function of the identified MYC N-terminal acidic clusters. First, we confirmed similar expression levels in HEK293T cells of MYC-WT and MYC-TAM by immunoblot analysis (Figure 4A) and flow cytometry (Figure 4B). HALO pull-down followed by immunoblot analysis also confirmed that MYC-TAM retains the ability to bind MYC’s obligate partner MAX (Figure 4C).

**Figure 4:**
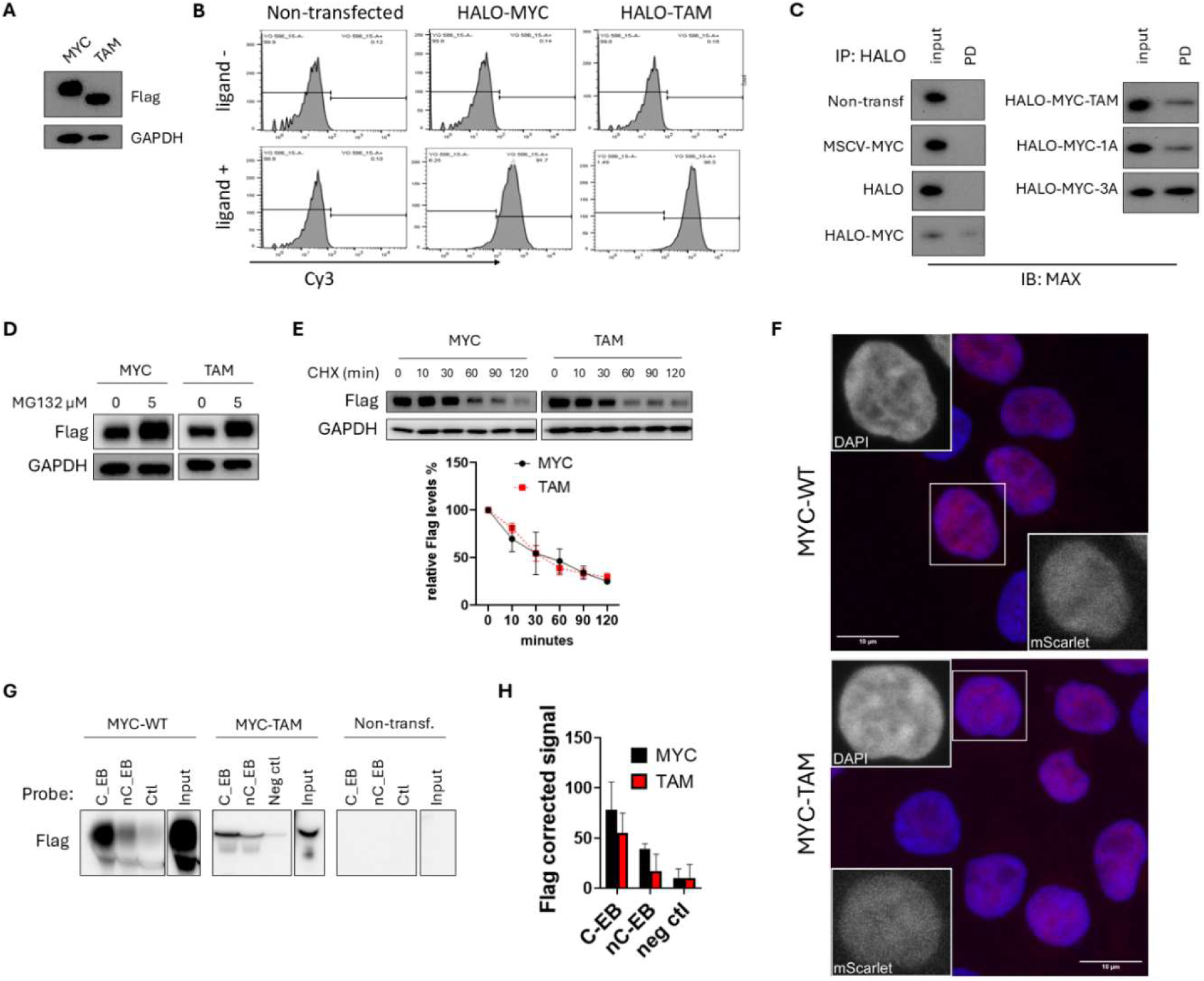
N-terminal acidic clusters do not regulate MYC protein stability, DNA-binding nor sub-cellular localization. A) HEK293T cells were transduced to stably express 3xFlag-MYC-WT or 3xFlag-MYC-TAM and nuclear extracts were immunoblotted for Flag expression. GAPDH was used as loading control. B) Flow cytometry analysis of 4188 cells expressing ectopic HALO-MYC-WT or HALO-MYC-TAM and incubated with the fluorescent HALO-specific TMR/Cy3 direct ligand. C) Pull down using HALO-link beads in nuclear extracts from 4188 cells expressing MYC-WT, HALO-MYC-WT, HALO-MYC-TAM, HALO-MYC-with neutralising mutants of acidic patch 1 (D>A; HALO-MYC1A) and 3 (D/E>A; HALO-MYC3A) or HALO control. Inputs and pull-down (PD) eluates were immunoblotted and analysed for the presence of MAX. D) HEK293T cells were incubated with MG132 for 4 h and the accumulation of 3xFlag-MYC-WT and 3xFlag-MYC-TAM was assessed in whole cell extracts by immunoblot. E) Cycloheximide (CHX) chase assay performed in HEK293T cells expressing 3xFlag-MYC-WT or 3xFlag-MYC-TAM. Whole cell extracts were obtained after incubation with CHX 50µg/mL for 0, 10, 30, 60, 90 and 120 minutes and immunoblotted against Flag. Quantification of immunoblots was performed by ImageJFiji and GAPDH was used as loading control. Mean±SD (n=2-4). F) Fluorescence images of HEK293T cells expressing ectopic mScarlet-MYC or mScarlet-TAM (red). DAPI (blue) was used for nuclear staining. G) oligo pulldown and immunoblot for MYC-WT and MYC-TAM. dsDNA probes containing the CACGTG canonical E-box motif (C-EB), the non-canonical CACCTG motif (nC-EB) or non-specific sequence control (ctl) were incubated with nuclear extracts of cells expressing MYC-WT, MYC-TAM or non-transfected control. H) Quantification of the oligo pull-down. Each band signal is normalized to the corresponding input (n=2).

The stability of the MYC protein plays a major role in modulating its function. The ubiquitin-proteasome system is the main degradation pathway that controls the protein turnover, although degradation by calpain-mediated cleavage has been also described (43, 44). In cells treated with the proteasome inhibitor MG132, the accumulation of MYC-WT and MYC-TAM was similar (Figure 4D), demonstrating that, like MYC-WT, MYC-TAM is still subject to ubiquitin-proteasome-mediated degradation. We also compared the degradation kinetics between MYC-WT and MYC-TAM by cycloheximide (CHX) chase analysis and determined that there was no significant difference on the turnover of MYC-WT and MYC-TAM proteins (Figure 4E).

To examine whether the acidic patches of the MYC N-terminus are involved in modulating MYC’s subcellular localization, we analysed HEK293T cells expressing mScarlet-MYC-WT or mScarlet-MYC-TAM by confocal microscopy. Both proteins showed similar localization profiles with a clear enrichment in the nucleus concentrating on euchromatic regions of DNA defined by areas that stained less brightly by DAPI (Figure 4F).

To investigate if DNA binding may be modulated by the acidity of the MYC N-terminus, we developed an oligo-pull down analysis using biotinylated double stranded DNA probes containing the CACGTG canonical E-box MYC binding motif, the non-canonical CACCTG motif or non-specific sequences as control (Supplementary table 1). Our DNA-oligo pull down analysis revealed that MYC-WT and MYC-TAM specifically bind to both canonical and non-canonical E-boxes with a stronger binding detected on the former (Figure 4G, H), therefore confirming that the acidic clusters of the MYC N-terminus are not involved in regulating MYC-DNA binding. We conclude that the acidic patches identified on the MYC N-terminus are not involved in the MYC:MAX interaction, the modulation of the protein stability nor subcellular localization. Furthermore, the negative charges of this domain do not regulate the DNA binding capacity of MYC.

### The N-terminal acidic clusters of MYC interact with several histone acetyl-transferase complexes (HATs)

MYC activity is modulated by a series of protein-protein interactions with a wide number of cofactors, the majority of which involve the conserved MYC boxes (MBs)(45). We reasoned that the acidic patches identified on the MYC N-terminus may be involved in the binding between MYC and cofactors that are critical in regulating its oncogenic activity. To identify MYC binding partners, proximity-dependent biotin labelling coupled to mass spectrometry (BioID-MS) was conducted on cells expressing FlagBioID-MYC-WT and FlagBioID-MYC-TAM. Our proteomics analysis identified 99 proteins in the MYC-WT interactome, including previously described MYC cofactors such as MAX, EP400, TXNL1, EPC1 or MBTD1 (BioID-MYC-WT vs BioID, Log2FC>1.5, adj. p-val<0.05; Supplementary Table 2). Mutation of the acidic clusters in MYC-TAM led to a significant reduction of interaction in a total of 32 factors (Figure 5A). Gene Ontology (GO) analysis revealed that most of these cofactors are involved in histone modification and chromatin remodelling (Figure 5B), including subunits of histone acetyltransferase (HAT) complexes such as STAGA and NuA4, specifically TRRAP, GCN5 (KAT2A), and EP400 (Figure 5C). Mutation of the acidic clusters also disrupted the interaction between MYC and SWI/SNF complex subunits, including KAT5, SMARCE1, and ARID1B. Orthogonal pull-downs coupled to immunoblot analysis confirmed interactions between MYC and TRRAP (subunit of STAGA and NuA4), ARID1B (subunit of SWI/SNF), and KAT2B (subunit of CBP/p300) and further validated the essential role of MYC N-terminal acidic patches in regulating these protein-protein interactions (Figure 5D). Our results demonstrate that MYC’s N-terminal acidic patches mediate interactions with chromatin-modifying complexes.

**Figure 5:**
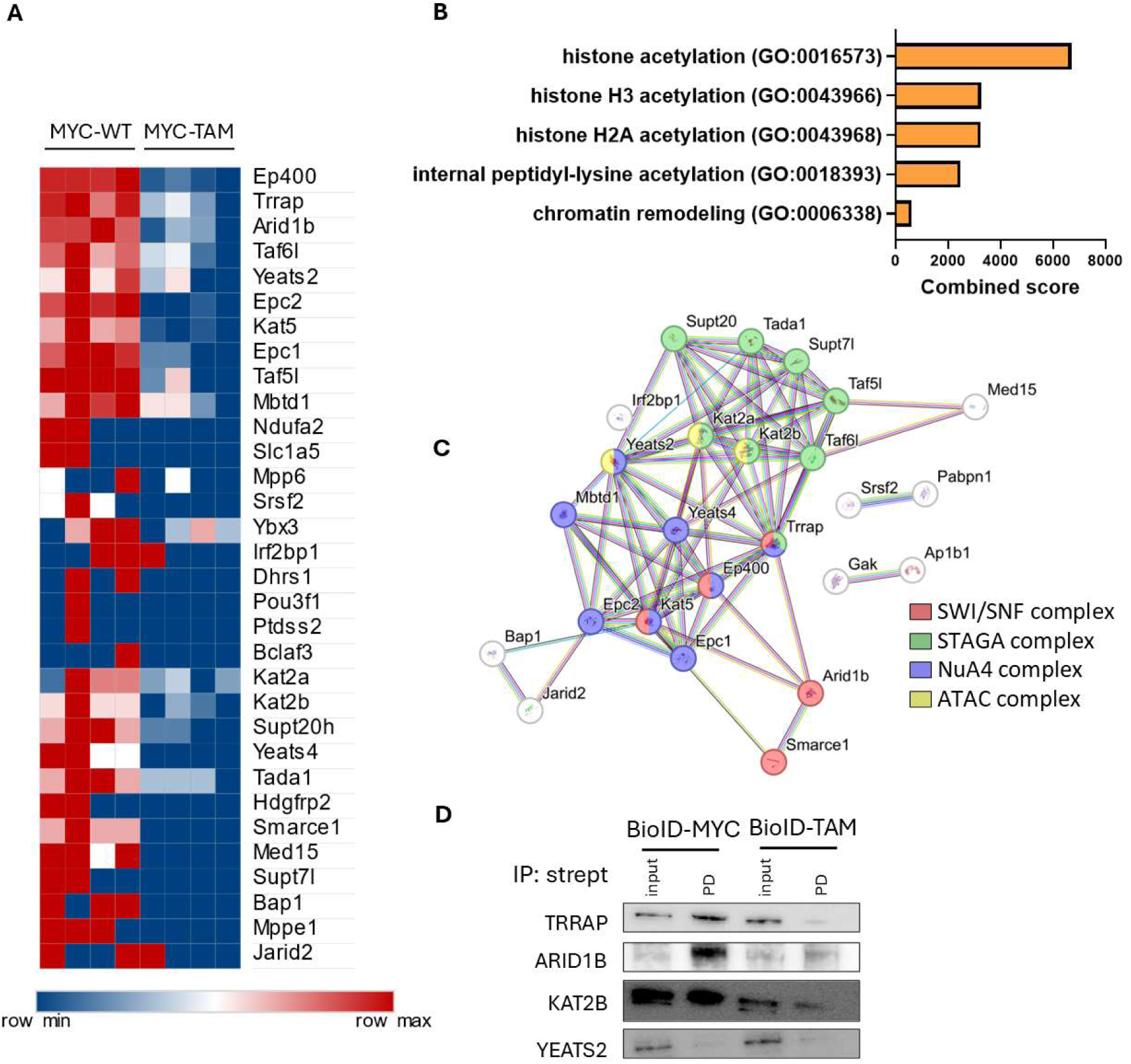
The N-terminal acidic patches of MYC interact with several HAT complexes. A) Heat map showing co-factors identified by BioID enriched in MYC-WT compared to MYC-TAM (log2FC<-1, FDR adj. p-value<0.05, n=4). In red, proteins with higher protein label-free quantification intensity. MYC-WT: cells expressing BioID-MYC-WT fusion protein, MYC-TAM: cells expressing BioID-MYC-TAM fusion protein. B) List of top 5 enriched GO terms (Adj p-value<0.05) ranked by combined score associated with the co-factors identified in the proximity-labelling proteomics analysis. C) STRING network protein-protein interaction diagram of the MYC cofactors associated with N-terminal acidic clusters. Colours indicate the components significantly associated (FDR Adj p-value<0.05) to each protein. D) Pull down using streptavidin beads in 4188 whole cell extracts expressing Flag-BioID control, Flag-BioID-MYC-WT or Flag-BioID-MYC-TAM. Inputs and pull-down (PD) eluates were immunoblotted and analysed for the presence of TRRAP, ARID1B, KAT2B and Yeats2 (n=3-4).

### MYC N-terminal acidic patches regulate the interaction with TRRAP

The high-throughput screening cancer dependency map DepMap ((https://depmap.org/portal) identifies potential RNAi co-dependency genes that are crucial for tumour survival, metastasis, and recurrence in more than 700 different cancer cell lines. Hypergeometric enrichment analysis showed a significant overlap (p-val= 2.11 × 10^-5^) between the MYC interactors identified by our proximity labelling proteomics and genes with dependency scores in DepMap corelated with MYC (Pearson coefficients ≥ 0.3). In addition, some of the MYC binding partners appear in the top 10% of ranked MYC co-dependent genes including EP400, KAT2B, or KAT5 (supplementary table 3) supporting a potential mechanistic link between interaction and co-essentiality. The protein TRRAP was ranked as the top protein coding gene even above the obligate MYC partner MAX indicating that the MYC:TRRAP interaction is indispensable for MYC oncogenic activity in a wide range of cancer cell lines (Supplementary figure 10A). This high co-dependency between MYC and TRRAP was confirmed in T-ALL by knocking-down TRRAP following lentiviral transduction of MYC-dependent Jurkat. TRRAP knock-down (Supplementary figure 10B) was associated with a decrease in cell viability compared to control safe-harbour AAVS1 (46) at 5 days post-transduction (Supplementary figure 10C), supporting the importance of TRRAP in modulating MYC activity in a T-ALL cell context.

TRRAP is a co-factor subunit of multiple HAT complexes including STAGA (GCN5/PCAF) and NuA4 (TIP60/p400) (47) (48). It has been described as an essential cofactor for MYC-dependent transformation by linking chromatin modification with MYC oncogenic activity (49). The interaction between MYC and TRRAP has been interrogated previously and the recruitment of TRRAP has been attributed to different regions of MYC such as MBII (50), the whole 1-110 N-terminal segment of MYC (51) or the M1, M2 and M3 subdomains distributed along the MYC N-terminus (52). However, our BioID MS data suggests that the group of acidic clusters located at the MYC N-terminus are essential for the interaction between MYC and TRRAP. Conversely within TRRAP, the region 2033-2088 has been previously described as the minimal domain involved in the interaction with MYC (50, 53). The role of MYC N-terminal acidic clusters in the MYC-TRRAP interaction was further validated in living cells using nano-bioluminescence resonance energy transfer (nanoBRET) based on the energy transfer from the bioluminescent donor Luciferase to the fluorescently labelled energy acceptor HALOtag. Briefly, the Luciferase-tagged TRRAP peptide (1899–2401) which contains the 2033-2088 minimal region involved in the interaction with MYC, was co-expressed with MYC-WT-HALO or MYC-TAM-HALO HEK293T cells. Upon addition of HALOTag-618 Ligand, the signal emitted from the excited fluorescent HALOtag was quantified (Figure 6A). Cells expressing MYC-WT showed an increased signal compared to those expressing MYC-TAM (p<0.05)(Figure 6B). To further prove the link between the N-terminal acidic clusters and TRRAP, we measured TRRAP recruitment into MYC condensates using a cell-based condensate system (54). CFP-MYC-WT was tethered to LacO array in U2OS cells expressing ectopic yellow fluorescent protein (YFP)-TRRAP_1997-2135_ fusion peptide, that contains the minimal region 2033-2088 (Figure 6C). YFP-TRRAP_1997-2135_ was enriched in tethered CFP-MYC-WT condensates and its recruitment was significantly decreased in the CFP-MYC-TAM condensates (p<0.001) (Figure 6D). These results suggest that the acidic patches of the MYC N-terminus are crucial for the interaction of MYC and TRRAP and for the recruitment of the latter to transcriptional condensates.

**Figure 6:**
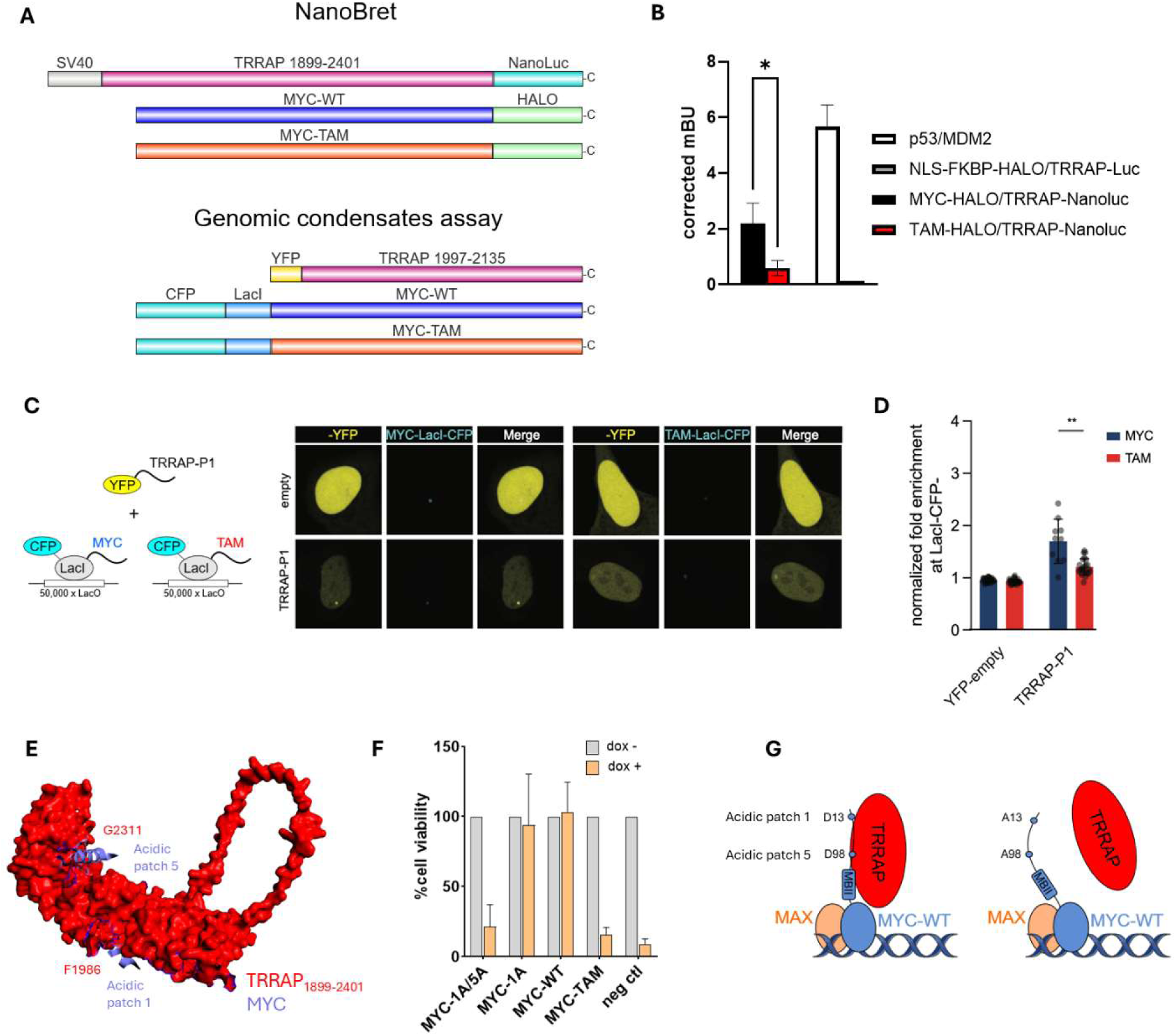
MYC N-terminal acidic patches regulate the interaction with TRRAP. A) Schematics of the constructs used for the validation of the identified protein-protein interactions: HALO pull-down, NanoBRET and genomic condensates assays. B) NanoBRET measurements of HEK293T cells co-transfected with MYC-WT-HALO/TRRAP_1899-2401_- NanoLuc or MYC-TAM-HALO/TRRAP_1899-2401_-NanoLuc. Co-transfection with P53-HALO/MDM2-NanoLuc was used as positive control and NLS-FKBP-HALO/TRRAP-Luc as negative control. Unpaired student t-test was performed. Mean and SD are showed. P<0.05*. C) Fluorescence images of ectopically expressed YFP-TRRAP_1997-2135_ in U2OS cells co- transfected with CFP-LacI-MYC-WT or CFP-LacI-MYC-TAM fusion constructs. D) Quantification of the relative YFP-TRRAP_1997-2135_ signal intensity in the CFP-LacI-MYC-WT or CFP-LacI-MYC-TAM foci. P values are from a Welch’s t-test. E) Representative model from AlphaFold3 of the N-terminal acidic patches 1 and 3 (blue) and the residues 1899-2401 of TRRAP (red). F) Cell viability of 4188 cells expressing the single mutant MYC-1A (acidic patch 1 mutated to alanine), MYC double mutant MYC-1A/5A (acidic patches 1 and 5 mutated to alanine), Flag-MYC-WT, Flag-MYC-TAM or parental 4188 (neg ctl) and incubated for 72h with doxycycline (20ng/mL). G) The negative charge of acidic clusters 1 and 5 within the intrinsically-disordered MYC N-terminus are predominant in regulating MYC:TRRAP binding, even in the presence of MBII.

Structural characterization of MYC is challenging due to its high conformational flexibility. To understand which N-terminal acidic patch(es) play a more relevant role in the interaction with TRRAP, we generated an AlphaFold3 predictive structure model of the complex (Figure 6E). Within the acidic patch 1 and acidic patch 5, residues D17 and D97 show a moderate interaction (<8Å between Cα-Cα) with TRRAP residues T2198 and F1986. Other residues within the patches exhibit smaller interaction distances such as MYC-Y16 with TRRAP-F1986 (5.1 Å), MYC-V19 with TRRAP-P1988 (5.1 Å), or MYC-L99 with TRRAP-G2311 (5.2 Å). Our prediction also shows the presence of a helical region containing MYC residues 96-105 (pLDDT score>50), a structure that is lost upon charge neutralization of MYC N-terminal acidic patches. Our functional analysis on 4188 expressing ectopic MYC-double mutant for acidic patch 1/5 in the presence of doxycycline revealed a loss of function in a magnitude similar to that observed for the MYC-TAM (Figure 6F), confirming the critical role of residues within the acidic clusters 1 and 5 in modulating the oncogenic function of MYC.

Overall, our results support a model in which MYC N-terminal acidic patches are required for the interaction between MYC and TRRAP even in the presence of an intact MBII domain (Figure 6G). These residues are also important in modulating the interaction with HATs and other chromatin modifier complexes modulating MYC transcriptional activity. Ultimately, MYC-dependent cell transformation and T-ALL tumour initiation are regulated by the acidic patches of MYC N-terminus.

## DISCUSSION

For decades, MYC has been appreciated as a major oncogene across multiple cancer subtypes, yet its therapeutic targeting has proven highly challenging. Most studies investigating the N-terminus have focused solely on the functional relevance of different MBs, despite the fact that many areas outside these domains remain highly conserved across diverse species (55–59). In this study, through site-saturation mutagenesis structure-function screening, we identify negatively charged clusters within MYC’s intrinsically disordered N-terminal domain as essential for its biological activity. We validate their functional relevance across multiple model systems both in vitro and in vivo, and provide new mechanistic insight into their role in mediating interactions with specific chromatin-modifying complexes.

We demonstrate that N-terminal acidic patches do not modulate the DNA-binding capacity of MYC, nor its interaction with MAX, but instead mediate an interaction with several HAT complexes. Of these, we propose that the interaction with the scaffold protein TRRAP is one of the most crucial, since it is required for MYC-dependent oncogenic transformation (49, 60) and has a strong co-dependency with MYC across hundreds of cancer cell line models in the DepMap dataset. TRRAP is a member of the phosphoinositide 3 kinase-related kinases (PIKKs), but is considered a pseudokinase due to the lack of a catalytic domain (61). TRRAP acts as a pivotal scaffold subunit required for the recruitment of histone acetyl transferase multiprotein complexes such as STAGA (SPT3–TAF9–GCN5 Acetyltransferase complex) or 60kDa Tat Interacting Protein (TIP60, also named NuA4) (53, 62). Original mapping analyses indicated that the MBI and MBII regions are essential for cell transformation and for MYC-TRRAP interaction, with MBII being the predominant interaction domain (63). The N-terminal regions M2 (within MBI) and M3 (residues 100-106) have been also implicated in the interaction with TRRAP (52). Although certain residues of the MYC N-terminal acidic patches (i.e. E54 or E101) are situated within the M2/M3 domain, mutations of the corresponding individual acidic clusters were not associated with a loss of MYC activity in our tet-off 4188 cell model. In contrast, our results implicate two highly conserved acidic clusters (acidic patches 1 and 5) located outside MBI and MBII as previously unappreciated key contributors to the MYC–TRRAP interface.

Our findings suggest a more distributed and multivalent binding mechanism that resonates with the conceptual framework proposed by Paul Sigler, who described “acidic blobs and negative noodles” as functional elements within disordered regions of transcription factors (64). These negatively charged segments are proposed to mediate electrostatic interactions with basic surfaces on other proteins, suggesting charge distribution, rather than fixed structure, plays a critical role at the interaction interface of such IDRs. Indeed, structural studies, such as those investigating the mediator subunit Med15 have shown acidic activation domains interact at a ‘fuzzy’ protein interface that is dynamic and unstructured (65).

Acidic patches in IDRs, such as those in BRD4 and OCT4, have been associated with the formation of phase separated transcriptional condensates (66, 67). Due to the challenges in characterising such condensates in live cells, we did not investigate the role that acidic patches might play in phase separation of the MYC IDR. However, we were able to show that the IDR of eIF4G2, which has been shown to form phase separated condensates when fused to other transcription factor complexes (39), was unable to rescue the activity of MYC-TAM. This would be consistent with the notion that acidic patches are critical to formation of a functional activation complex through mediation of electrostatic interactions, rather than phase separation per se. Further work would be required to conclude this definitively.

In summary, we have identified new acidic domains critical for MYC oncogenic function that modulates its global transcriptional activity via its interaction with several HAT complexes, in part through the interaction with the adaptor protein TRRAP. We demonstrate that the interaction between MYC and TRRAP is not solely dependent on MBII, but instead orchestrated by a broader interaction surface. The conservation and functional indispensability of these acidic patches across models underscore their evolutionary and biological importance. Lastly, and perhaps most importantly, the identification of a distinct, functionally essential subdomain within MYC uncovers new vulnerabilities that are likely relevant for therapeutic targeting.

## METHODS

### Cell culture

Low passage HEK-293T cells were cultured in DMEM supplemented with 10% FBS, l-glutamine, and penicillin/streptomycin. The murine T-cell line 4188 was cultured in RPMI-1640 supplemented with 10% FBS and 1% penicillin/streptomycin. Doxycycline (Merk) was used at 20 ng/mL. Protease inhibitor MG132 (Cambridge Bioscience) was used at 5 µM. Cycloheximide (CHX) (Merck) was used at 50 µg/mL for the indicated time points. MCF10A cells were a kind gift from Dr Bart Vanhaesebroeck from UCL Cancer Institute (UK). MCF10A cells were cultured in Dulbacco’s modified Eagle’s medium (DMEM)/F12 (Sigma) supplemented with 5% donor horse serum (Sigma), 20 ng/ml of epidermal growth factor (EGF, Preprotech), 0.5 mg/ml hydrocortisone (Sigma), 100 ng/ml cholera toxin (Sigma), 10 mg/ml insulin (Sigma), and 100 units/ml penicillin and 100 mg/ml streptomycin. SHEP-21N cells were a kind gift from Prof. Louis Chesler from Institute of Cancer Research (ICR). Low passage SHEP-21N cells were cultured in DMEM supplemented with 10% FBS, l-glutamine, and penicillin/streptomycin.

### Site-saturation mutagenesis screening

The MYC mutant library containing 7,140 single amino acid variants of mouse MYC covering positions 5-354, and C-terminal positions 357, 366, 382, 413, 420, 427 and 434 was generated and cloned into an MSCV retroviral backbone by Twist bioscience using proprietary technology. The library also contained a negative control consisting of residues 1-25 of MYC. Each mutated position was associated with a specific barcode at 3’. Retroviral transduction was conducted as indicated and 4188 cells were infected aiming for a MOI≤0.3 to ensure the integration of only one MYC variant per cell. To achieve this, dilutions of the retroviral supernatant were used to transduce ≈100*10^6^ cells that were then expanded for 2 days and selected with puromycin (2µg/mL) for 3 days. Cell viability was analysed using CellTiter-Glo and only those cultures with a cell viability <30% and thus ensuring ≥1000 cells per variant were expanded and selected for screening. Selected cells were split into equal fractions (16*10^6^ cells/fraction) and incubated with doxycycline (20ng/mL) for 96h. Cell viability was analysed again by CellTiter-Glo and cultures showing <10% reduction after doxycycline treatment where further processed. Genomic DNA (gDNA) was extracted using QIAMP DNA blood maxi kit (QIAGEN #51192) and processed for hybrid capture using the Twist Custom Panel Hybridization Capture of DNA Libraries kit (Twist bioscience #101057) according to manufacturer’s instructions. Briefly, gDNA was enzymatically fragmented, and ends were repaired and dA-tailed (#101058). Tailed DNA fragments were then ligated with indexed adapters specific for each condition (dox off, dox on)(Twist Bioscience #101059 and #100255). Indexed gDNA libraries were pre-captured PCR amplified and purified (Twist bioscience #101059) and amplified indexed libraries were pooled (750 ng each). Hybrid capture was performed using a custom-made pool of biotinylated probes (14x120bp each, with 4bp overlap) that capture the mouse gene DNA sequence (gene ID 17869, that translates for protein NP_034979). Hybridized targets were bound to streptavidin beads and enriched libraries were obtained following the post-capture PCR amplification and purification of targets. Repeated QC checks were performed along the library prep process after each PCR amplification using D5000 high-sensitivity tapescreen (Agilent #5067-5592 and #5067-5593) and QUBIT HS dsDNA (ThermoFischer #Q32851).

Samples were prepared according to the Illumina® Nextera XT protocol for NGS analysis. Libraries were indexed with unique i7/i5 index pairs and after the limited-cycle PCR step, reactions were combined and purified using AMPure XP beads (Beckman Coulter, #A63880. Samples were analysed using the NextSeq™ 550 Sequencing System (Illumina®) (250PE).

NextSeq™ 550 Sequencing data was processed using a sequence caller for the analysis of sgRNA representation in CRISPR screenings (68) that was adapted to count the number of reads for each barcode. To determine the degree of enrichment, first we excluded those barcodes that showed <100 reads in the control (dox off) condition and computed the log2 fold change (log2FC) for the remaining barcodes. We assumed >50% of sites did not show a significant effect (supported by our cell viability data). We determined the median and standard deviation of the log2FC values and these were subtracted in order to compute the centered log2FC normalized to library size. Two-sided p-values were determined and corrected the p-values for multiple testing were obtained using the Benjamini–Hochberg method.

### SHEP-21N proliferation assay

SHEP21N cells were seeded on 96-well plates (2000 cells/well) and counted on days 1, 2, 3,4 and 7 using a hemacytometer. Cell doublings (CD) per hour were obtained using the formula (Nj-Ni)/ln2)/72h where Nj or Ni is the cell number at different time point Tj or Ti (Tj>Ti) during the cell growth log phase (days 4 to 7). Doubling time (DT) was determined by dividing the time interval (Tj>Ti) by the CD (69).

### Soft agar colony assay

The human mammary epithelial cell line MCF-10A is a widely used in vitro model for studying normal breast cell function and transformation that exhibits an increased anchorage-independent proliferation when overexpressing ectopic MYC (70). MCF10A cells were cultured in 6-well plates. Wells were first covered with an 0.6 % agar layer mixed with DMEM/F12 10%HS and, once solidified, a middle layer of 0.3 % agar mixed with 10% HS DMEM/F12 was added mixed with 5,000 cells per well.

SHEP21N neuroblastoma cells expressing tetracycline (tet) regulatable N-MYC were also used as model to evaluate MYC-dependent transformation in soft agar colony formation assay (70). SHEP21N cells were incubated overnight on 1µg/mL doxycycline to turn off endogenous expression of N-MYC and cultured 6-well plates. Wells containing solid 0.6 % agar layer mixed with 10%FBS DMEM were covered with a middle layer of 0.3 % agar mixed with 10% FBS DMEM containing 1µg/mL doxycycline or DMSO. Each well contained 5,000 cells.

Every 3-4 days, 300uL of the corresponding culture media (10% HS DMEM/F12 for MCF10A and 10%FBS DMEM containing doxycycline or DMSO for SHEP21N) were added into each well to prevent agar drying.

Cultures (n=4-8) were incubated for ≈14 days and stained in 0.005% crystal violet for colony counting. Cultures were imaged in a Bio-Rad ChemiDoc MP imaging system and colonies were counted on ImageJ FIJI.

### Retroviral transduction

Retrovirus were generated as previously described (71) by co-transfecting packaging plasmids VsVg (2 µg) and pMD.MLV (4 µg) along with constructs of interest. Plasmids were mixed on 30 µL of Genejuice (Merk) and 470 µL of OPTIMEM (Gibco) and used for transfection of HEK293T cells at 50% confluence in DMEM medium (10%FBS, 1% pen/strep). On the following day, DMEM medium was replaced by RPMI (10%FBS, 1% pen/strep) and the virus containing supernatant was collected on day 2 and day 3 after cell infection, pooled together and filtered through 0.45 µm filter pore size. Mouse 4188 cells (5*10^5^-1*10^6^ cells/well) were transduced by spinoculation (2500 rpm, 1.5 h at 37 °C) using the corresponding retroviral suspensions and 8 µg polybrene. After 4 h of incubation, the medium was replaced by fresh RPMI and cells incubated at 37 oC for 72h. Puromycin selection was conducted for 72h and cell viability was assessed using CellTiter Glo.

### Constructs generation

Acidic mutations on each cluster were introduced by Q5 site directed mutagenesis using an MSCVpuroMYC WT plasmid as template. A gBlock gene fragment containing the TAM mutations at the N-terminus of MYC flanked by XhoI and SbfI-HF restriction sites was digested and cloned into a MSCVpuro-MYC WT plasmid backbone. All primer sequences used can be found at “Q5 site directed mutagenesis general protocol”. The reverse acidic mutant constructs were generated following the same approach with specific primers for the E to D and D to E residue substitutions.

To generate the HALO-TEV-tagged version of each variant, the HALO-TEV tag on the 5’ end of the acidic mutants was added by PCR amplification using the forward primer “TATACTCGAGATTTCCGGCGAGCCAACCACTGAGGATCTGTACTTTCAGAGCGAT” and reverse primer “TATAGAATTCTTATGCACCAGAGTTTCGAAGCTGT”, using the corresponding acidic variant as template. The resulting product contains the C-terminal end of HALO, the TEV protease target sequence and the whole MYC protein sequence with each corresponding mutation, all flanked with XhoI and EcoRI-HF restriction sites. The PCR products were gel purified (QIAquick gel purification kit, QIAGEN), restriction enzyme digested, T4 ligated (NEB) into a compatibly linearized MSCVpuro-HALO plasmid.

The MYC acidic variants were PCR amplified using the forward primer 5’- TATACTCGAGATGCCCCTCAACGTGAACTTCACC-3’ and reverse primer 5’- TATAGTCGACGAATTCTTATGCACCAGAGTTTCGAAGCTGT-3’ and subsequently digested with XhoI/HincII (NEB). Digestion products where gel purified and cloned into a linearized pBABE-FlagBioID2 plasmid backbone.

For NanoBret analysis, MYC-WT and MYC-TAM were cloned into a halotag7-pfc14k-cmv-flexi-vector-eu113047 backbone (Promega) in frame with c-terminal HALO-tag whereas. The TRRAP_1899–2401_ region was cloned from a gBlock into nluc-cmv-neo-flexi-vector-pfc32k- complete-sequence-kf811456 (Promega) in frame with C-terminal NanoLuc. Both inserts were cloned by AsiSI and Eco53ki restriction digestion. The constructs NanoLuc-MDM2 (Promega) and P53-Halotag (Promega) fusion vectors were used as positive control whereas a construct containing NLS-BFP-FKBP-HA-HALOtag was used in combination with TRRAP-NanoLuc as negative control.

All constructs were transfected into Stellar (Takara) competent cells and were confirmed by Sanger sequencing.

### Amino acid enrichment and charge analyses

Amino acid frequency analysis was analysed using ProtParam tool from Expasy (https://web.expasy.org/protparam) (72). Net charge per residue for MYC was computed using the localCIDER tool (73).

### BioID experiments

Transfected 4188 cells expressing Flag-BirA* (BioID), Flag-BirA*-MYC-WT (BioID-MYC-WT), Flag-BirA*-MYC-TAM or control untransfected (120*10^6^cells per condition) were incubated in complete RPMI (10%FBS, 1% pen/strep) and 50µM biotin (SigmaAldrich) for 16 hours. Samples were all analysed in 4 biological replicates. Cells were washed 3 times in PBS and nuclear extracts were collected (NE-PER, Thermo). Final extracts were supplemented with cOmplete protease inhibitor cocktail (Roche) and 125U benzonase (SigmaAldrich). Pull-downs of biotinylated proteins were performed using the streptavidin-Sepharose beads (GE) previously derivatized according to the protocol published elsewhere (74). After the Sepharose-beads derivatization, these were washed in equilibration buffer (20mM HEPES, pH 7.5, 420 mM NaCl, 1.5 mM MgCl2, 0.2 mM EDTA). A total of 250uL of beads were incubated with 3 mg of total protein at a concentration of 0.5mg/mL with rotation overnight at 4 °C. Beads were then washed with 1mL x 4 with 50mM Tris-Cl, pH7.4, 8M urea and resuspended in 50uL of 5 mM biotin in 50mM ammonium bicarbonate (pH8). Beads were washed 5 times with 50mM TEAB and once with 50mM TEAB and urea 6M with a 2 min incubation. Urea was diluted to 2M with TEAB 50mM and pH adjusted to 8.5-9. Trypsin was diluted 1/10 in TEAB 50mM, 20μL were added into the solution containing the beads and incubated 2h at 37°C. Additional 20μL of trypsin 1/10 were added and incubated overnight at 37°C. After digestion, peptides were collected and pH adjusted to 2-3 using 10%TFA. C18 columns (17-170µg) were equilibrated with 100% ACN once and three times with 0.1% TFA. Samples were loaded into columns and washed four times with 0.1% TFA and 5% ACN. Finally, samples were eluted by adding 3x75µL of 0.1% TFA and 50% ACN and lyophilized using a Speedvac centrifuge (2000rpm, 4C, 3-3.5h).

### Mass spectrometry

nLC-MS/MS was performed on a Q Exactive Orbitrap Plus interfaced to a NANOSPRAY FLEX ion source and coupled to an Easy-nLC 1200 (Thermo Scientific). Five percent of each sample was loaded as 5 µL injections. Peptides were separated on a 27 cm fused silica emitter, 75 μm diameter, packed in-house with Reprosil-Pur 200 C18-AQ, 2.4 μm resin (Dr. Maisch) over 120 min using a linear gradient of 95:5 to 70:30 buffer A:B (buffer A: 0.1% formic acid in water; buffer B: 80% acetonitrile/0.1% formic acid), at a flow rate of 250 nL/min. Peptides were ionised by electrospray ionisation using 1.9 kV applied immediately prior to the analytical column via a microtee built into the nanospray source with the ion transfer tube heated to 320°C and the S-lens set to 60%. Precursor ions were measured in a data-dependent mode in the orbitrap analyser at a resolution of 70,000 and a target value of 3e6 ions. The ten most intense ions from each MS1 scan were isolated, fragmented in the HCD cell, and measured in the orbitrap at a resolution of 17,500.

### Protein identification and relative quantification

Raw data were analysed with MaxQuant (75) version 1.6.17 where they were searched against the mouse SwissProt database (http://www.uniprot.org/, downloaded 01/09/2021) using default settings. Carbamidomethylation of cysteines was set as fixed modification, and oxidation of methionines and acetylation at protein N-termini were set as variable modifications. Enzyme specificity was set to trypsin with maximally 2 missed cleavages allowed. To ensure high confidence identifications, PSMs, peptides, and proteins were filtered at a less than 1% false discovery rate (FDR). Label-free quantification in MaxQuant was used with LFQ minimum ratio count set to 2 with ‘FastLFQ’ (LFQ minimum number of neighbours = 3, and LFQ average number of neighbours = 6) and ‘Skip normalisation’ selected. In Advanced identifications, ‘Second peptides’ and the ‘match between runs’ feature were selected. For statistical protein quantification analysis, the ‘proteinGroups.txt’ and ‘evidence.txt’ output files from MaxQuant were loaded into the MSstats quantification statistical framework package (76) (version 3.14.0) run through RStudio (version 1.1.456, R version 4.0). Contaminants and reverse sequences were removed and data were log2 transformed, and a linear mixed-effects model was fitted to the data. As a large difference between the conditions was expected, data was not normalised. The group comparison function was employed to test for differential abundance between conditions. p-values were adjusted to control the FDR using the Benjamini-Hochberg procedure (77). The MYC-WT interactome was determined by proteins with a log2FC<-1, FDR adj p-value<0.05 in BioID-MYC vs Control comparison. From these, proteins identified in the BioID-MYC condition in at least one replicate were compared to BioID-TAM and those with log2FC<-1, FDR adj p-value<0.05 were selected for network and Gene Ontology analyses. Network analysis: physical interaction data was downloaded from the STRING database v10 (78).

### Cell viability assays

Cells were seeded (10,000 cells/well) in 96-well white plates for luminescence assays (CulturePlateTM, PerkinElmer) and treated with 20 ng/mL of doxycycline for 72/96 hours. Cell viability was determined using CellTiter-Glo Luminescent Viability Assay (Promega) as per manufacturer’s instructions, and luminescence was measured using Varioskan LUX Microplate Reader (Thermo Fisher Scientific).

### HALO TMR Direct ligand incubation and flow cytometry analysis

The HALO TMR Direct ligand 555Ex/585Em (Promega, catalog. G2991) was used to confirm protein expression by flow cytometry as indicated in manufacturer’s instructions. Briefly, transfected cells were incubated with 0.1 µM TMR Direct ligand for 1 hour at 37°C, washed with PBS and analysed by flow cytometry in a FACS canto (BD Biosciences).

### HALO pull-down

Nuclear extracts were obtained (Thermo Scientific, #78835) from cells expressing different HALO-tagged MYC proteins, supplemented with protease inhibitor (Thermo Scientific, # 78430) and quantified using BCA Protein Assay kit (Pierce, #23225). HALOlink resin (Promega, #G9410) was washed in 100mM Tris-HCl (pH 7.5), 150mM NaCl, 1mg/mL BSA, 0.05% IGEPAL® CA-630, mixed with samples containing a total of 0.5 mg total protein (0.5µg/µL, adjusted in TBS (150mM NaCl, 50mM Tris-HCl (pH 7.5)) and incubated overnight at 4°C. One percent of each sample was kept as input control. Samples were washed, resuspended in TEV buffer (50mM Tris-HCl (pH 8), 0.5 mM EDTA, 1mM DTT) containing TEV protease and incubated for 1.5 hours at room temperature. was performed. Eluates and inputs were further analysed by Western Blot.

### Western Blot

Samples were resolved in 4-12% Bis-Tris NuPAGE gels (Thermo Scientific, # NP0335PK2) at 80V for 10 minutes followed by 120V for 1.5 hours and then transferred to a 0.22-μm PVDF membrane (Maine Manufacturing, #1215037) preactivated with 100% methanol using the Mini-trans blot system (Bio-Rad, #1703930) at 100V for 2 hours at 4^0^C. Membranes were blocked either with 5% BSA (Sigma, # A9418) or milk (Sigma, #70166) in PBS 0.1% Tween 20 (PBS-T) and incubated overnight with a primary antibody in 2.5% BSA or milk at 4°C. Membranes were washed 3 times in PBS-T and probed probing with the corresponding horseradish peroxidase (HRP)–conjugated secondary antibody in 2.5% BSA or milk PBS-T for 1 hour. Membranes were washed 3 times and developed on a Chemidoc MP reader (Bio-Rad, #17001402) using Immobilon Western Chemiluminescent HRP substrate (Sigma, #42029053).

The following primary antibodies were used in this study: c-Myc (D84C12) Rabbit mAb (Cell Signalling Technologies, catalog. 5605), MAX polyclonal rabbit Antibody (Thermo Fisher Scientific, catalog. PA5-79637), TRRAP) rabbit polyclonal (Invitrogen, catalog. PA5-78246), FLAG mouse monoclonal (Merk, catalog. F1804), ARID1B mouse monoclonal (Santa Cruz, catalog. sc-32762), KAT2B rabbit monoclonal (Thermo Fisher Scientific, catalog. MA511186), GAPDH rabbit monoclonal (Cell Signalling, catalog. 5174S). Secondary antibodies used include Anti-mouse IgG, HRP-linked antibody (Cell Signalling, catalog. 7076P2) and Anti-rabbit IgG, HRP-linked Antibody (Cell Signalling, catalog. 7074).

### Oligo pull-down

Biotinylated oligonucleotide probes were generated as previously described (79). Oligonucleotide probes were bound to paramagnetic streptavidin beads (Dynabeads MyOne Streptavidin C1, Invitrogen) before being incubated with nuclear protein extracts in PBB buffer (150 mM NaCl, 50 mM Tris-HCl pH 7.5, 5 mM MgCl2, 0.5% IGEPAL CA-630 (Sigma)) in the presence of sheared salmon sperm DNA (Ambion) for 2 hours at 4°C on a rotating wheel. Three PBB buffer washes were performed and bound proteins were eluted in 2x Laemmli buffer and boiled for 5 minutes at 95°C. Eluates were analysed by immunoblot using anti-HA antibody (Cell Signalling). Blots are quantified using ImageJ. Background signal is subtracted from each band and corrected by the corresponding input.

### RNA-seq

Total RNA was extracted from 10,000,000 cells (4188 cell line) per sample using the RNeasy Mini Kit (Qiagen). mRNA was purified from total RNA using poly-T oligo-attached magnetic beads. Fragmentation was carried out using divalent cations under elevated temperature in First Strand Synthesis Reaction Buffer(5X). First strand cDNA was synthesized using random hexamer primer and M-MuLV Reverse Transcriptase (RNase H-). Second strand cDNA synthesis was subsequently performed using DNA Polymerase I and RNase H. Remaining overhangs were converted into blunt ends via exonuclease/polymerase activities. After adenylation of 3’ ends of DNA fragments, Adaptor with hairpin loop structure were ligated to prepare for hybridization. To select cDNA fragments of preferentially 370∼420 bp in length, the library fragments were purified with AMPure XP system (Beckman Coulter, Beverly, USA). Then PCR was performed with Phusion High-Fidelity DNA polymerase, Universal PCR primers and Index (X) Primer. At last, PCR products were purified (AMPure XP system) and library quality was assessed on the Agilent Bioanalyzer 2100 system. The clustering of the index-coded samples was performed on a cBot Cluster Generation System using TruSeq PE Cluster Kit v3-cBot-HS (Illumina) according to the manufacturer’s instructions. After cluster generation, the library preparations were sequenced on an Illumina Novaseq platform and 150 bp paired-end reads were generated. The data discussed in this publication have been deposited in NCBI’s Gene Expression Omnibus (Edgar et al., 2002) and are accessible through GEO Series accession number GSE291777 (https://www.ncbi.nlm.nih.gov/geo/query/acc.cgi?acc= GSE291777).

Raw data (raw reads) of fastq format were firstly processed through in-house perl scripts. In this step, clean data (clean reads) were obtained by removing reads containing adapter, reads containing poly-N and low quality reads from raw data. At the same time, Q20, Q30 and GC content the clean data were calculated. All the downstream analyses were based on the clean data with high quality.

Reference genome and gene model annotation files were downloaded from genome website directly. Index of the reference genome was built using Hisat2 v2.0.5 and paired-end clean reads were aligned to the reference genome using Hisat2 v2.0.5. The mapped reads of each sample were assembled by StringTie (v1.3.3b) (80) in a reference-based approach. FeatureCounts v1.5.0-p3 was used to count the reads numbers mapped to each gene and FPKM (Fragments Per Kilobase of transcript sequence per Millions base pairs sequenced) of each gene was calculated. Differential expression analysis of two conditions/groups (three biological replicates per condition) was performed using the DESeq2 R package (1.20.0). DESeq2 provide statistical routines for determining differential expression in digital gene expression data using a model based on the negative binomial distribution. The resulting P-values were adjusted using the Benjamini and Hochberg’s approach for controlling the false discovery rate. Genes with an adjusted P-value <=0.05 found by DESeq2 were assigned as differentially expressed.

Gene Set Enrichment Analysis v 4.2.3(GSEA) was conducted using genes ranked according to the degree of differential expression in the two samples and using both, predefined Gene Sets and own generated Gene Sets containing the 500 genes most upregulated or most downregulated (FDR<10^-99^) in non-transfected MYC-off 4188 cells.

### Genomic condensates assays and confocal imaging

The peptide YFP-TRRAP(1997–2135) was co-expressed with CFP-LacI-MYC-WT or CFP-LacI-MYC-TAM in U2OS cells containing the lac operon (LacO). Cells were plated on sterilized glass coverslips precoated with poly-L lysine in 6-well plates. Once adhered, cells were fixed with 3.7% PFA (Sigma) and permeabilized with 0.25% Triton X-100 (Merk). Those cells stained with antibodies were previously in blocking solution (PBS 5%BSA) and later blotted with the corresponding antibodies diluted in 0.5x blocking solution 0.1% saponin and stained with Hoetch (Merk). Cells that were not labelled with antibodies were directly stained with Hoetch (Merk). Coverslips were finally mounted on glass slides using ProLong Diamond mounting solution (Thermo). Images were collected using a Zeiss LSM 900 Confocal.

### DepMap data RNAi analysis

The DepMap (https://depmap.org/portal/) RNAi dataset consists of LOF screening results from 713 cell lines covering 17,309 genes in total. Co-dependency scores were obtained between MYC and the rest of the genes. We ranked each gene according to the obtained Pearson’s correlation coefficient when compared to MYC. We assessed enrichment of MYC interactors among MYC co-dependent genes using a hypergeometric test. The background set included 16,767 genes profiled in the DepMap. From this, we identified a subset of 4 genes showing Pearson correlation coefficients ≥ 0.3 with MYC dependency scores across cell lines. The set of MYC interactors consisted of 32 proteins identified through BioID-MS. Overlap between these 32 interactors and the 4 MYC co-dependent genes was tested using the hypergeometric distribution in R, with parameters N=16767, m=4, n=32, and k=2.

### TRRAP CRISPR knock out

TRRAP CRISPR sgRNA guides sgRNA1: 5’-CAGCATTCCATCATTCCGA-3’, sgRNA2: 5’- CCACTGGGGATCGTTCAGTG-3’, sgRNA3: 5’- CTTGATCCGCCACTATACGA-3’, and sgRNA4: 5’- TGGTGTCAAGACAATCACGT-3’ from the Brunello Knock-Out library (81) were cloned into a lentiCRISPRV2 plasmid backbone (Addgene # 244694). The safe-harbour AAVS1 sgRNA: 5’- AGCGGCTCCAATTCGGAAGT-3’ (46) was used as control. Lentiviral particles were following the same steps in the production of retroviral particles but using packaging plasmids VsVg and psPAX2. Jurkat cells were transduced (5*10^5^-1*10^6^ cells/constructs) as by spinoculation (2500 rpm, 1.5 h at 37 °C) using the corresponding retroviral suspensions and 8 µg polybrene. Cells were incubated for 48h followed by puromycin selection. Cell viability (CellTiter Glo) data and cell lysates were obtained at 72h after adding puro.

### Nanobret details

A total of 0.8*10^6^ HEK293T cells were transfected with 2µg of HaloTag® Fusion Vector DNA + 0.2µg of NanoLuc® Fusion Vector following the manufacturer’s protocol. Briefly, after mixing the corresponding plasmids combinations in Opti-MEM® I Reduced Serum Medium, no phenol red (ThermoFisher) and GeneJuice (Merk), plasmids were added into the cell cultures and incubated 20 hours at 37°C, 5% CO2 to allow protein expression. A total of 2,000 cells per well were seeded in a 96-well plate (CulturePlateTM, PerkinElmer) in the presence of HaloTag® NanoBRET™ 618 Ligand or DMSO vehicle (1nM) and incubated overnight at 37°C, 5% CO_2_. NanoBRET™ Nano-Glo® Substrate (Promega) was diluted 1/100 and added in Opti-MEM® I Reduced Serum Medium, no phenol red. Nanobret donor emission (460nm) and acceptor emission (618nm) were quantified using a Varioskan LUX Microplate Reader (Thermo Fisher Scientific).

### Protein disorder predictions

Disorder predictions were obtained from the corresponding FASTA sequence and using the models PONDR (https://pondr.com/) and ADOPT (82).

### Statistical analysis

All data are represented as mean ± SD. Statistical analysis was performed using GraphPad Prism 10 (GraphPad Software, La Jolla, CA, USA). Unpaired student t-test was performed when two groups were involved unless stated otherwise.

### Data availability

The mass spectrometry proteomics data have been deposited to the ProteomeXchange Consortium via the PRIDE (83) partner repository with the dataset identifier PXD048459 (Project Name: Identifying MYC interactors using BioID labelling, Project accession: PXD048459, Project DOI: Not applicable, Reviewer account details: Username, Password: FnrRoQHF). The RNA-seq data discussed in this publication have been deposited in NCBI’s Gene Expression Omnibus (84) and are accessible through GEO Series accession number GSE291777 (https://www.ncbi.nlm.nih.gov/geo/query/acc.cgi?acc=GSE291777).

## ACKNOWLEDGEMENTS

We would like to thank all the members of the Mansour lab for their support and advice that helped make this project possible. We are grateful for the insightful scientific discussions with Dr. Alejandro Gutierrez and Dr. Alex Kentsis regarding this project. We thank Alex McLatchie for his assistance with next-generation sequencing experiments. We also thank Deni Kappei for his advice on the oligo pull-down approach and Elisabeth Henderson for her advice on the NanoBRET construct design and analysis. We would like to acknowledge the support of the Proteomics Research, the Flow Cytometry and Imaging TTP. The current work was funded by CRUK Pioneer Award (C61186/A26238) and Therapeutic Discovery (DRCPLT-Nov20100001) programmes. The Devices & Diagnostics TIN Pilot Data fund partially supported the study through the MRC CiC6 award. V. L and Y.L were funded by John Goldman Fellowships from Leukaemia UK (2022/JGF/003 and 2024/JGF/002, respectively). M.R.M is funded through a Great Ormond Street Children’s Charity professorship. D.O.C was funded by CRUK/Children with Cancer (grant DRCPGM∖100066).

## AUTHORS CONTRIBUTIONS

V.L and M.R.M conceptualized the study. V.L designed the wet lab experiments. Bioinformatic and transcriptomic analyses were performed by V.L, D.O.C, J.D, K.F and L.W. Site-saturation mutagenesis screening, mutagenesis experiments, functional analyses, flow cytometry, cycloheximide chase, proteasome inhibition, protein pull-down, NanoBRET analyses and soft-agar colony formation assays were performed and analysed by V.L. A.T cloned the constructs for BioID-MS analyses and S.S and A.B. performed the proteomics analyses and assisted with validation. V.L, T.R.D.S and F.A performed western blots, generated and maintained cell lines. V.L and Y.L obtained and analysed confocal images. H.N, I.S and D.H generated the constructs, performed transcriptional condensates recruitment experiments and analysed the data generated. A.T.L and S.H performed and analysed the experiments with zebrafish. AlphaFold modelling was performed by S.B, C.F and V.L. M.R.M supervised the work and experiment design. V.L and M.R.M drafted the manuscript. All authors contributed to critical discussions and approved the final manuscript.

---

**Supplementary table 1.**
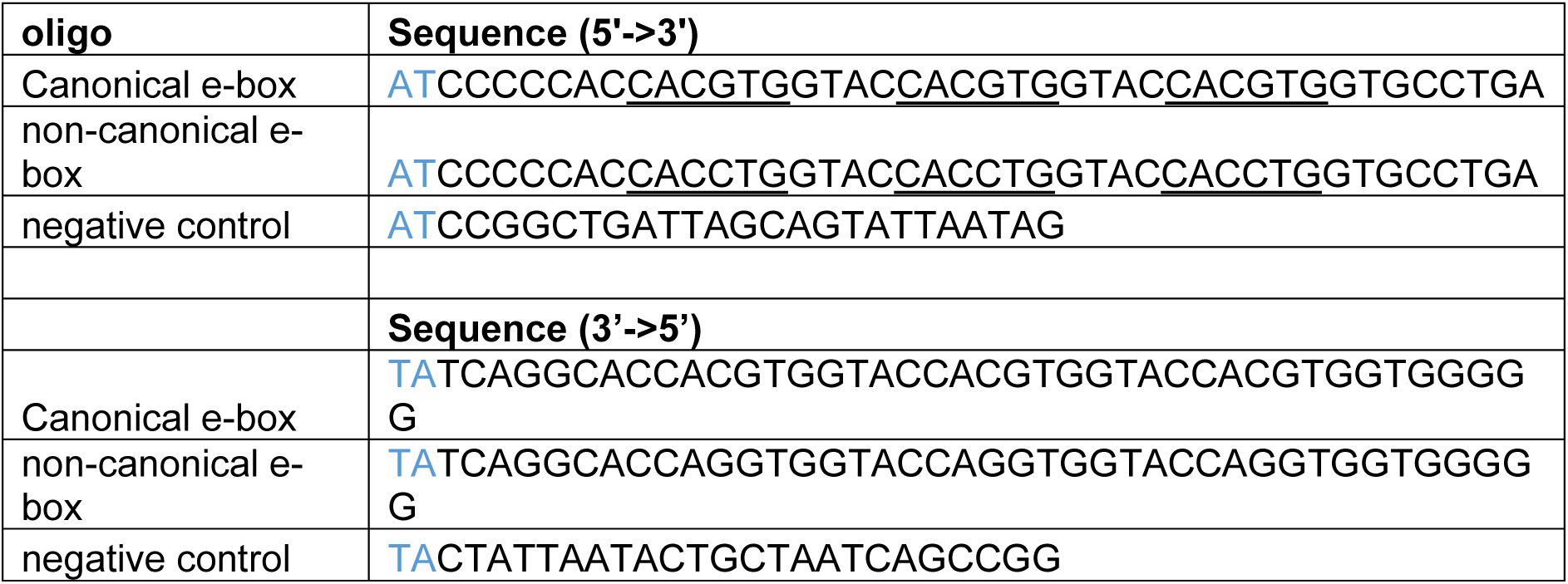
Oligos designed for concatemerization. Motifs contained are underlined. Forward sequences contain overhang (blue).

**Supplementary table 2.**
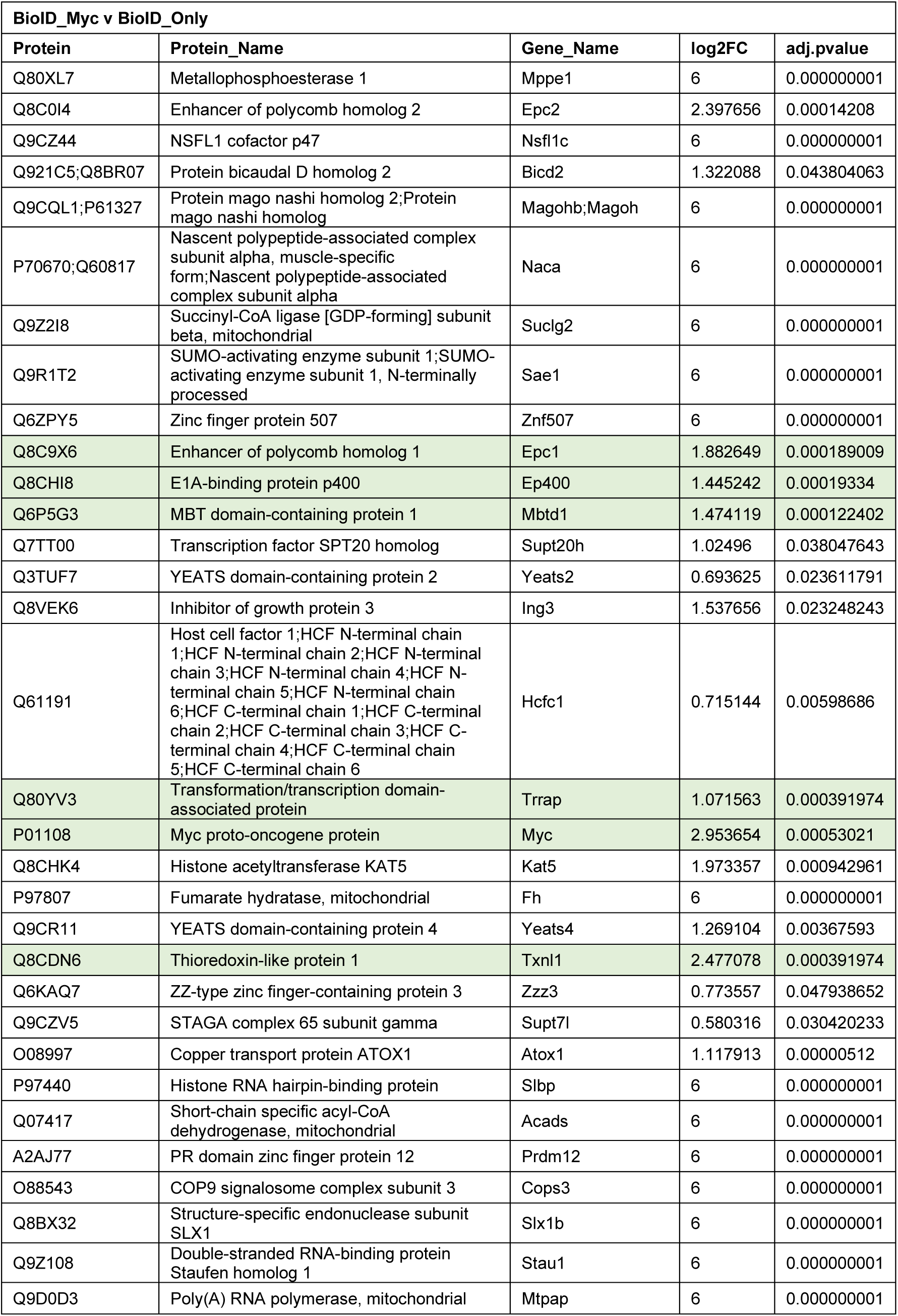
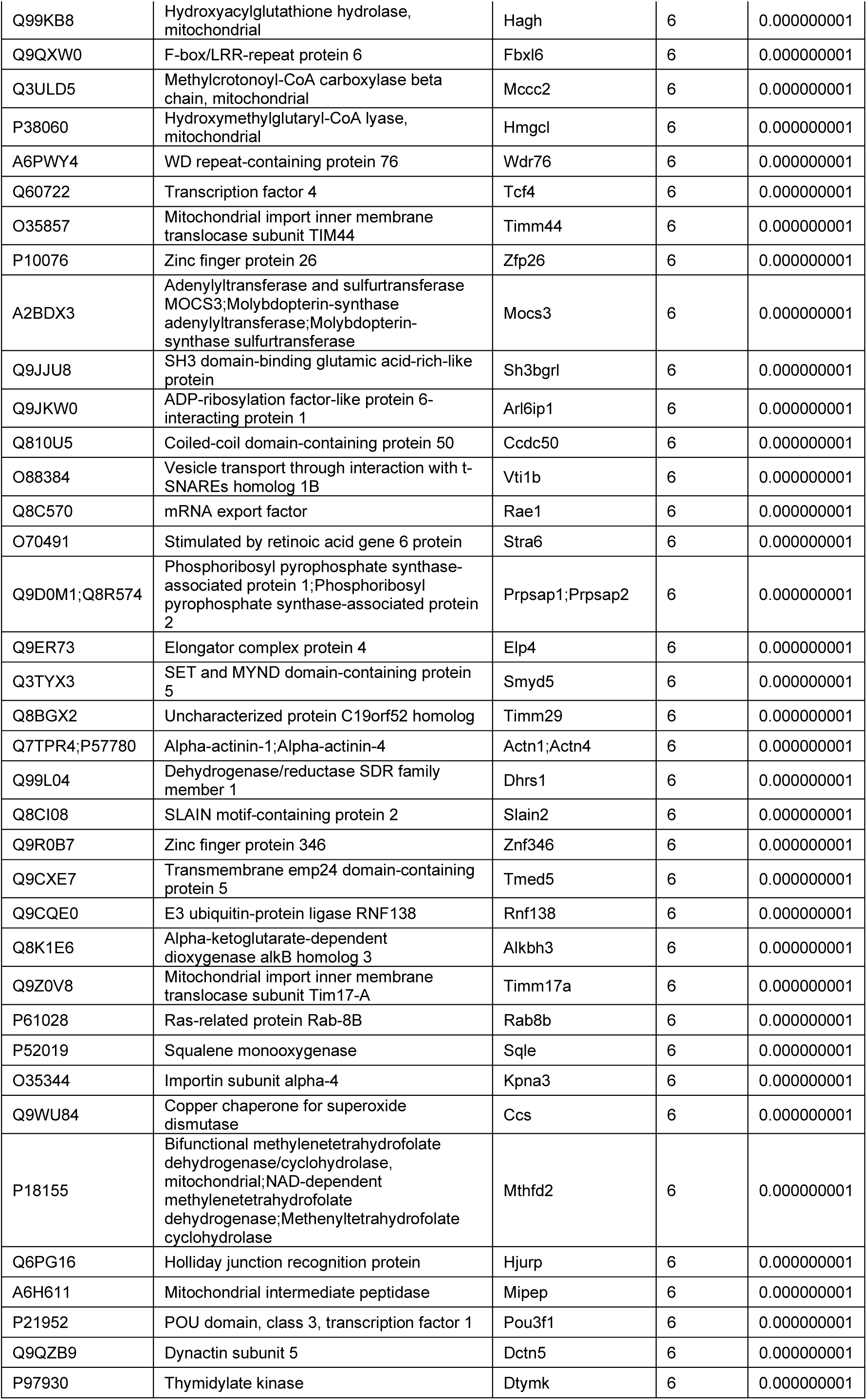
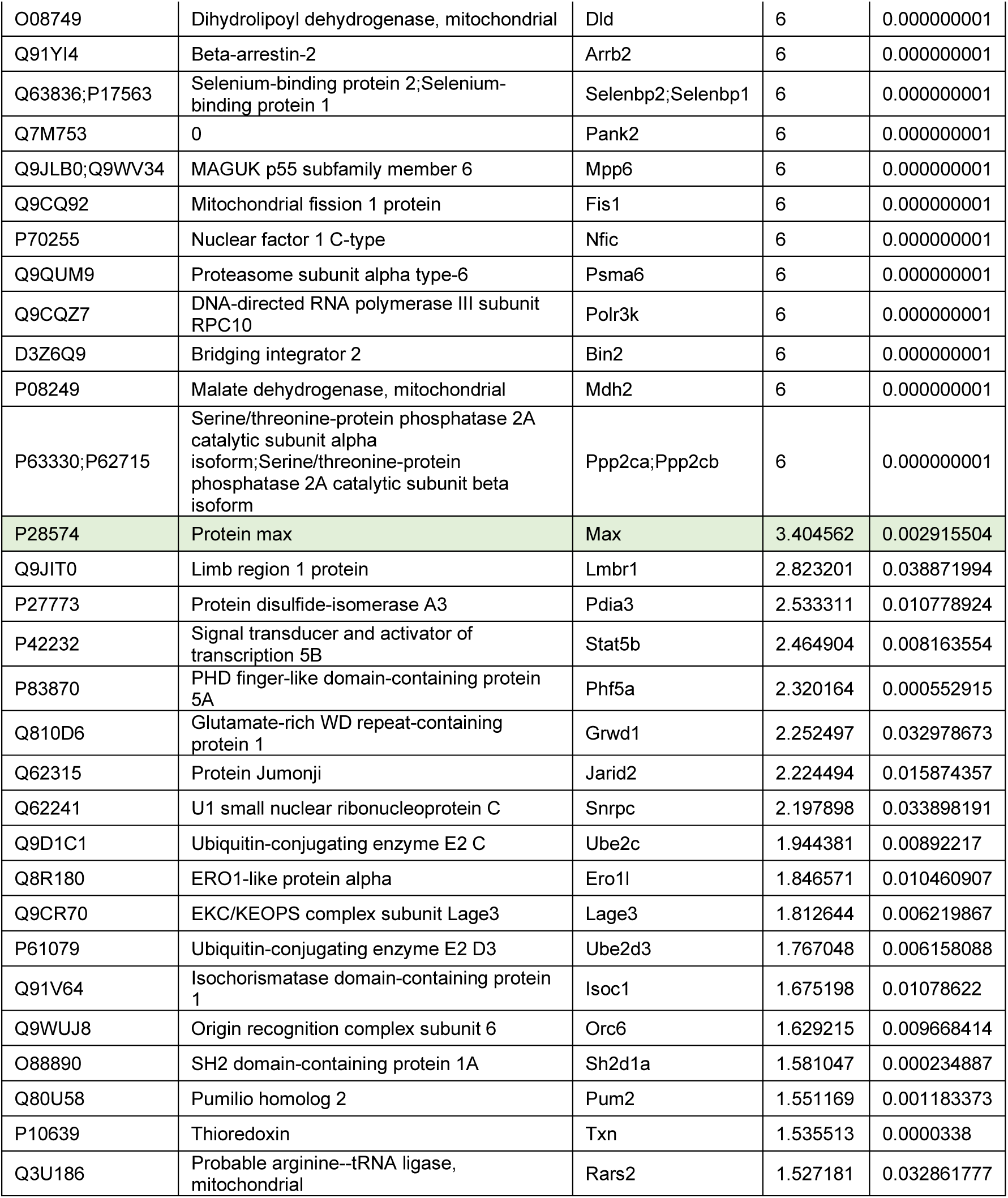
Protein enriched in the FlagBioID-MYC-WT interactome compared to FlagBioID control (logFC>1.5 and adj. p-val<0.05). Highlighted in green previously described MYC interactors MAX, EP400, TXNL1, EPC1 or MBTD1 and TRRAP.

**Supplementary table 3.**
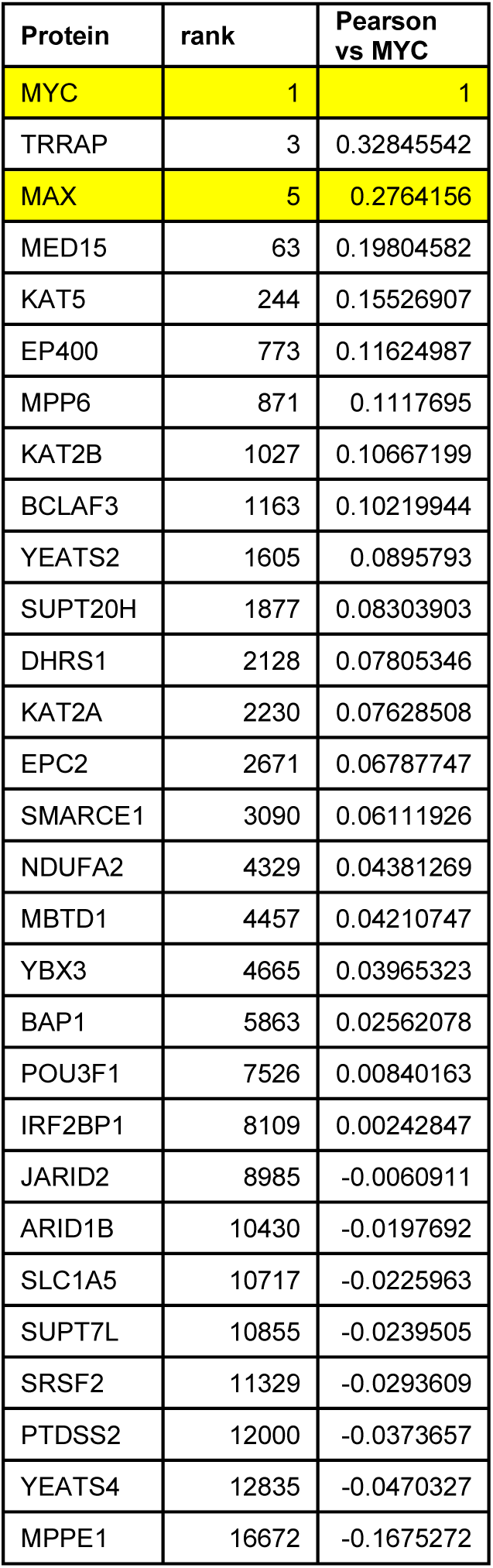
The co-dependency score between 16,767 proteins and MYC was obtained for 712 cell lines from DepMap portal https://depmap.org/portal/. This table indicates the co-dependency score of the MYC interactors identified that depend on MYC’s N-terminal acidic patches for binding. MYC and MAX have been added into the list.

| Protein | rank | Pearson<br>vs MYC |
| --- | --- | --- |
| MYC | 1 | 1 |
| TRRAP | 3 | 0.32845542 |
| MAX | 5 | 0.2764156 |
| MED15 | 63 | 0.19804582 |
| KAT5 | 244 | 0.15526907 |
| EP400 | 773 | 0.11624987 |
| MPP6 | 871 | 0.1117695 |
| KAT2B | 1027 | 0.10667199 |
| BCLAF3 | 1163 | 0.10219944 |
| YEATS2 | 1605 | 0.0895793 |
| SUPT20H | 1877 | 0.08303903 |
| DHRS1 | 2128 | 0.07805346 |
| KAT2A | 2230 | 0.07628508 |
| EPC2 | 2671 | 0.06787747 |
| SMARCE1 | 3090 | 0.06111926 |
| NDUFA2 | 4329 | 0.04381269 |
| MBTD1 | 4457 | 0.04210747 |
| YBX3 | 4665 | 0.03965323 |
| BAP1 | 5863 | 0.02562078 |
| POU3F1 | 7526 | 0.00840163 |
| IRF2BP1 | 8109 | 0.00242847 |
| JARID2 | 8985 | -0.0060911 |
| ARID1B | 10430 | -0.0197692 |
| SLC1A5 | 10717 | -0.0225963 |
| SUPT7L | 10855 | -0.0239505 |
| SRSF2 | 11329 | -0.0293609 |
| PTDSS2 | 12000 | -0.0373657 |
| YEATS4 | 12835 | -0.0470327 |
| MPPE1 | 16672 | -0.1675272 |

## Supplementary figures

**Supplementary figure 1:**
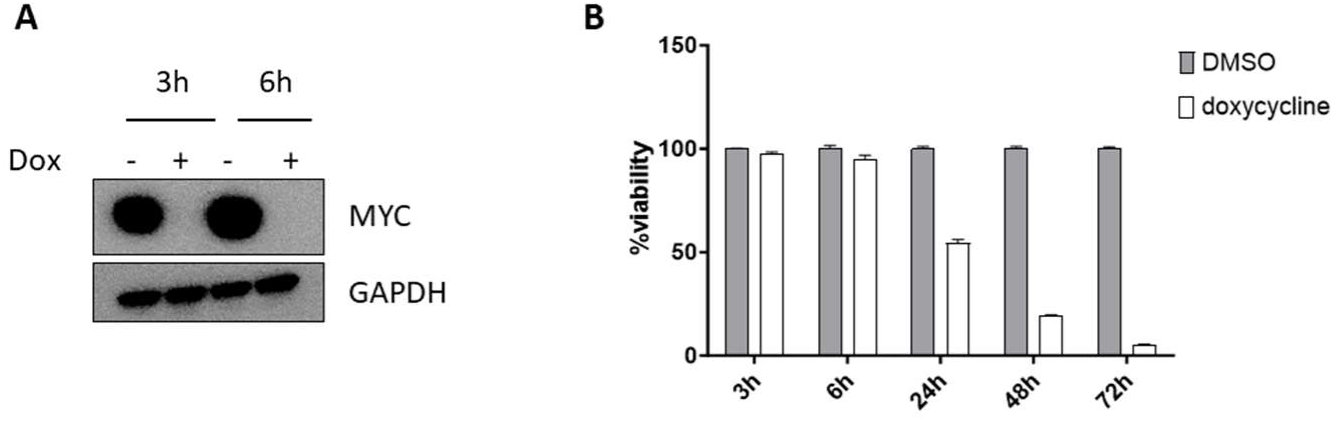
A) Immunoblot showing the expression of endogenous MYC in 4188 cells whole lysate after incubation with doxycycline (20ng/mL) or DMSO at different time points. GAPDH was used as loading control. B) Cell viability of 4188 measured by Cell Titter Glo after incubation with doxycycline (20ng/mL) or DMSO at different time points.

**Supplementary figure 2:**
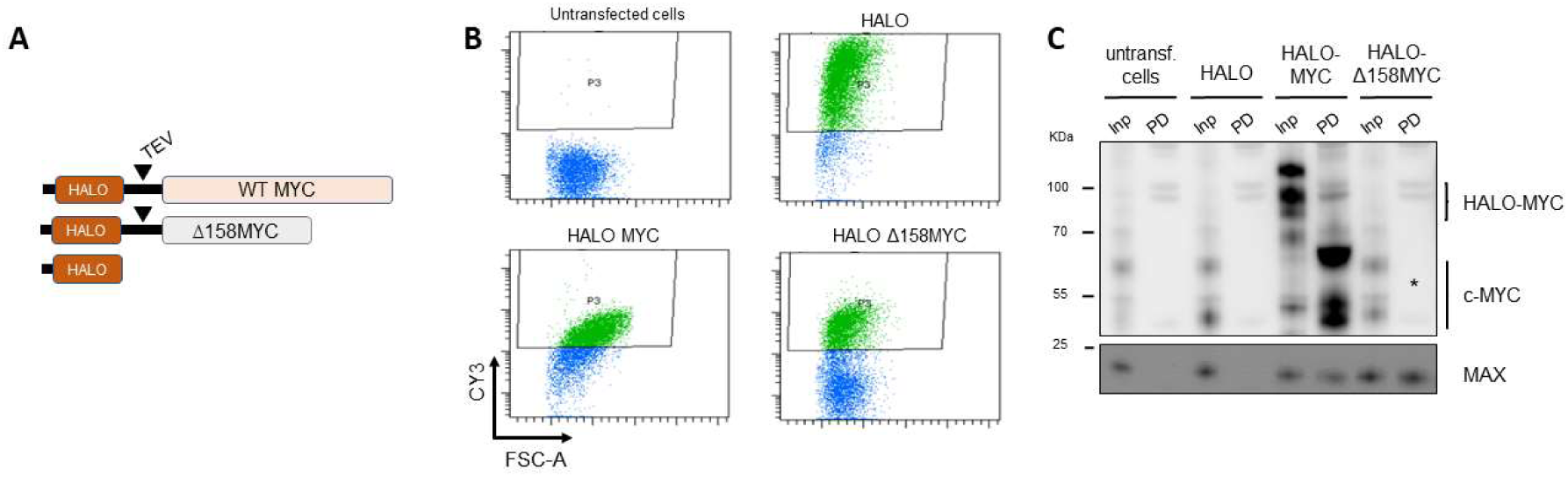
A) Diagram of HALO-tagged constructs tested. MYC wild type (MYC-WT) MYC bearing the 1-158 deletion (Δ158MYC) were N-terminally tagged with the HALO moiety. HALO-tag alone was used as negative control. B) 4188 cells retrovirally transduced with HALO-MYC-WT, HALO-Δ158MYC or HALO were incubated with the HALO-specific TMR/Cy3 direct ligand and analysed by flow cytometry. All transduced cell cultures were positive for Cy3 confirming the expression of the corresponding fusion proteins. C) HALO-pull down and immunoblots using nuclear extracts from 4188 cells stably expressing HALO, HALO-MYC-WT and HALO-Δ158MYC. Western blot analysis the presence of MAX on the eluates of HALO-MYC-WT and HALO-Δ158MYC confirming that deletion of 1-158 does not prevent the MYC:MAX interaction. c-MYC indicates the bands corresponding to endogenous c-MYC detected in all inputs. The absence of Δ158MYC after HALO-pull down and TEV cleavage (*) results from the removal of the antibody-recognized epitope after deleting 1-158 portion of c-MYC.

**Supplementary figure 3:**
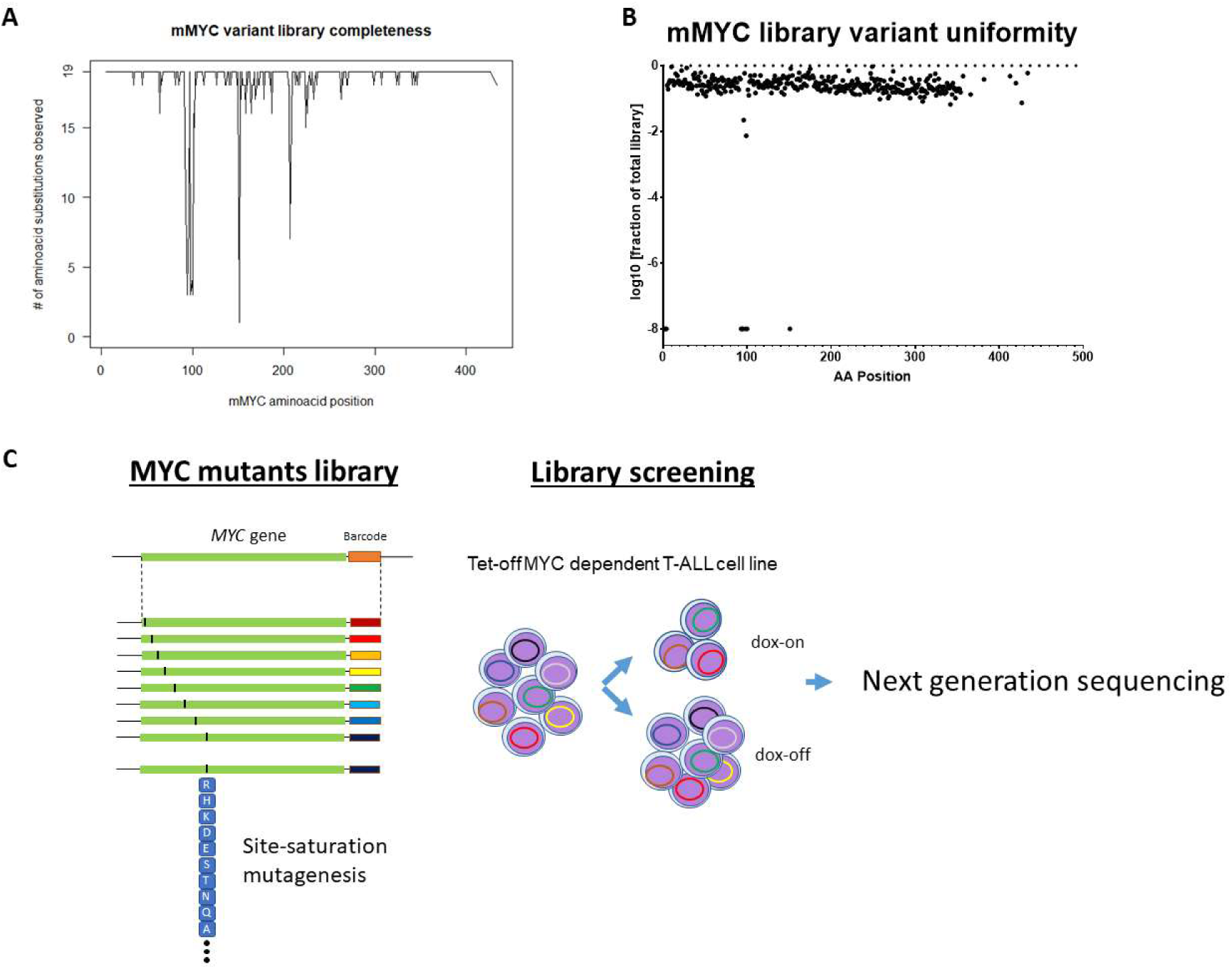
Site-saturation mutagenesis MYC library was generated by PCR incorporation of oligonucleotides carrying each variant modification. A) MYC variant library completeness plot showing the number of amino acid substitutions achieved per position. Site-saturation mutagenesis is achieved in one position when the wild type of amino acid residue is replaced by each of the other 19 amino acid variants (number of substitutions is 19). A site was considered to pass if >85% of the variants are detected. Each variant was deemed to be detected it reached a read count >20% of the theoretical expected value by next generation sequencing analysis. B) Dot plot showing the MYC library uniformity after transformation and expansion of E. coli competent cells. Uniformity was evaluated analysing the presence of each barcode by next generation sequencing. C) Schematic of the experimental workflow. A site-saturation MYC mutant library comprising 6783 MYC variants was introduced into T-ALL 4188 cells using an MOI<0.3 favouring the integration of a single variant in each cell. Library-transduced cells were treated with doxycycline (dox-on) or with DMSO (dox-off). By day 4, genomic DNA was collected, PCR amplified and analysed by next generation sequencing.

**Supplementary figure 4:**
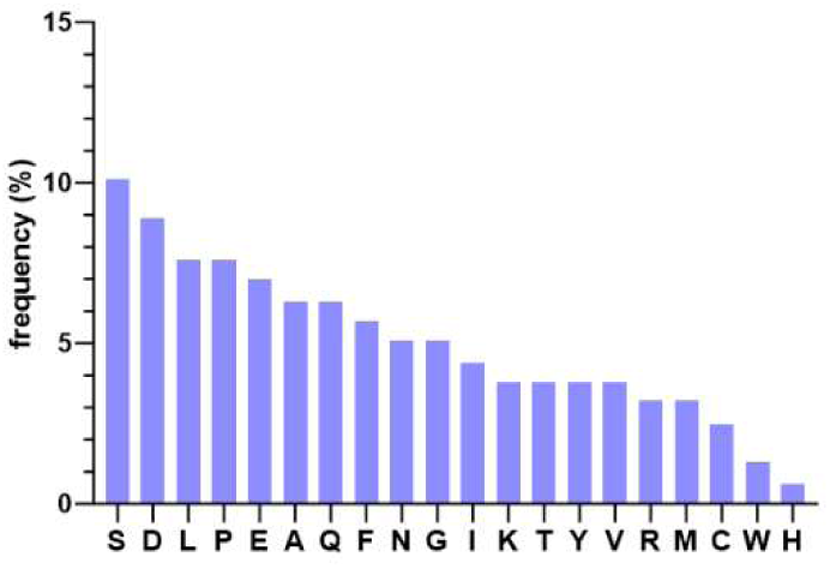
Relative amino-acids composition of the 1-158 N-terminal domain of MYC.

**Supplementary figure 5:**
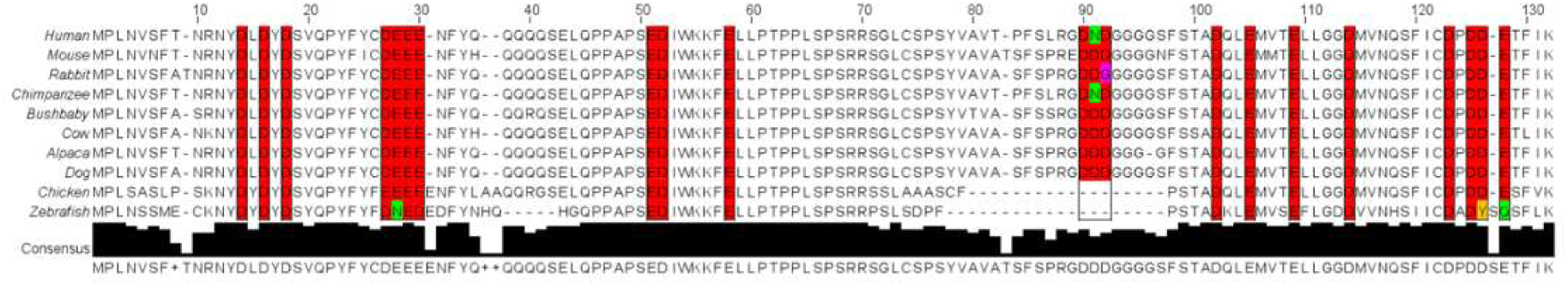
Sequence conservation of MYC protein positions 1-132 across species. Individual residues from the identified acidic clusters are highlighted. In red, those residues that preserve the acidic charge.

**Supplementary figure 6:**
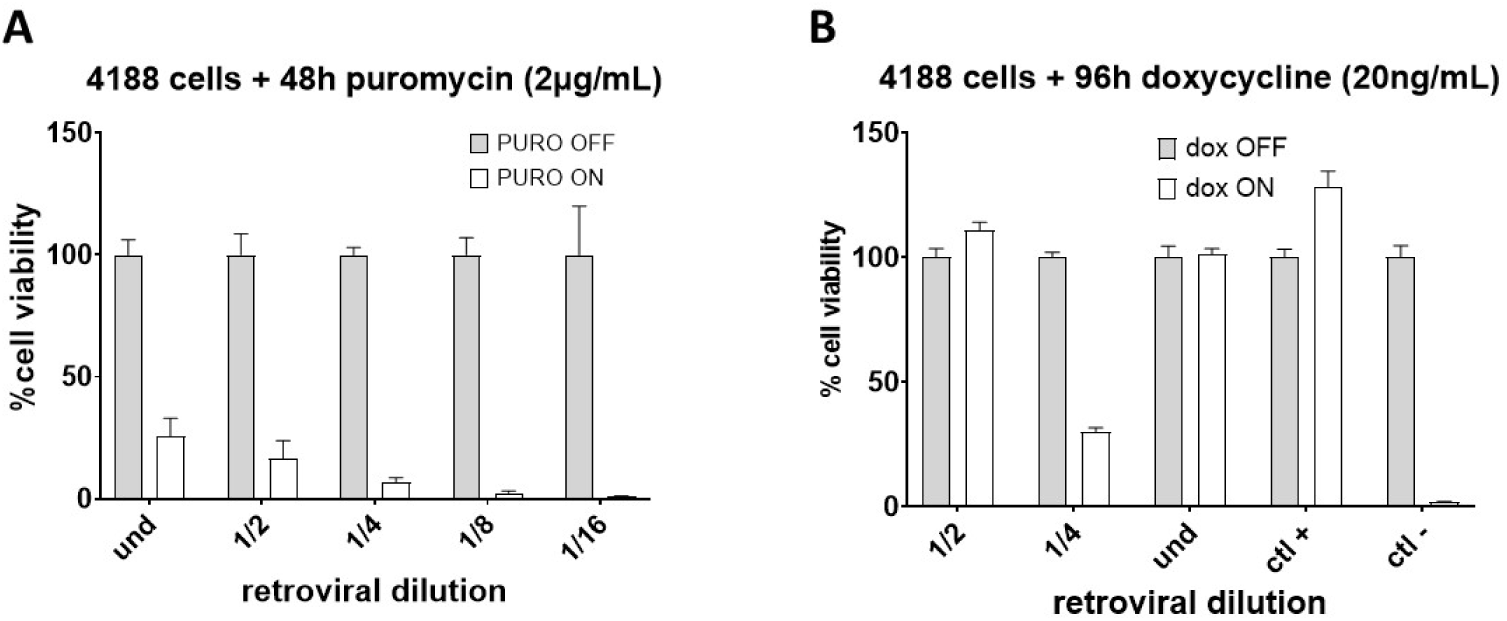
The rescue capacity of acidic mutants decreases with low MOI. 4188 cells were transduced using different dilutions of retrovirus MSCV-MYC-4A containing the alanine mutations of acidic patch 4. Cells were subsequently puromycin-selected and cell viability was measured. The retroviral dilution associated with a cell viability <30% post-puromycin was selected to test the rescue capacity of cell viability under doxycycline treatment. A) Bar graph shows viability of 4188 cells after selection with puromycin (2μg/mL) for 48h. B) Cell viability of 4188 cells after incubation with doxycycline (20ng/mL) for 96h and inhibition of endogenous MYC. Retroviral dilution ¼ exhibits cell viability ≈30%.

**Supplementary figure 7:**
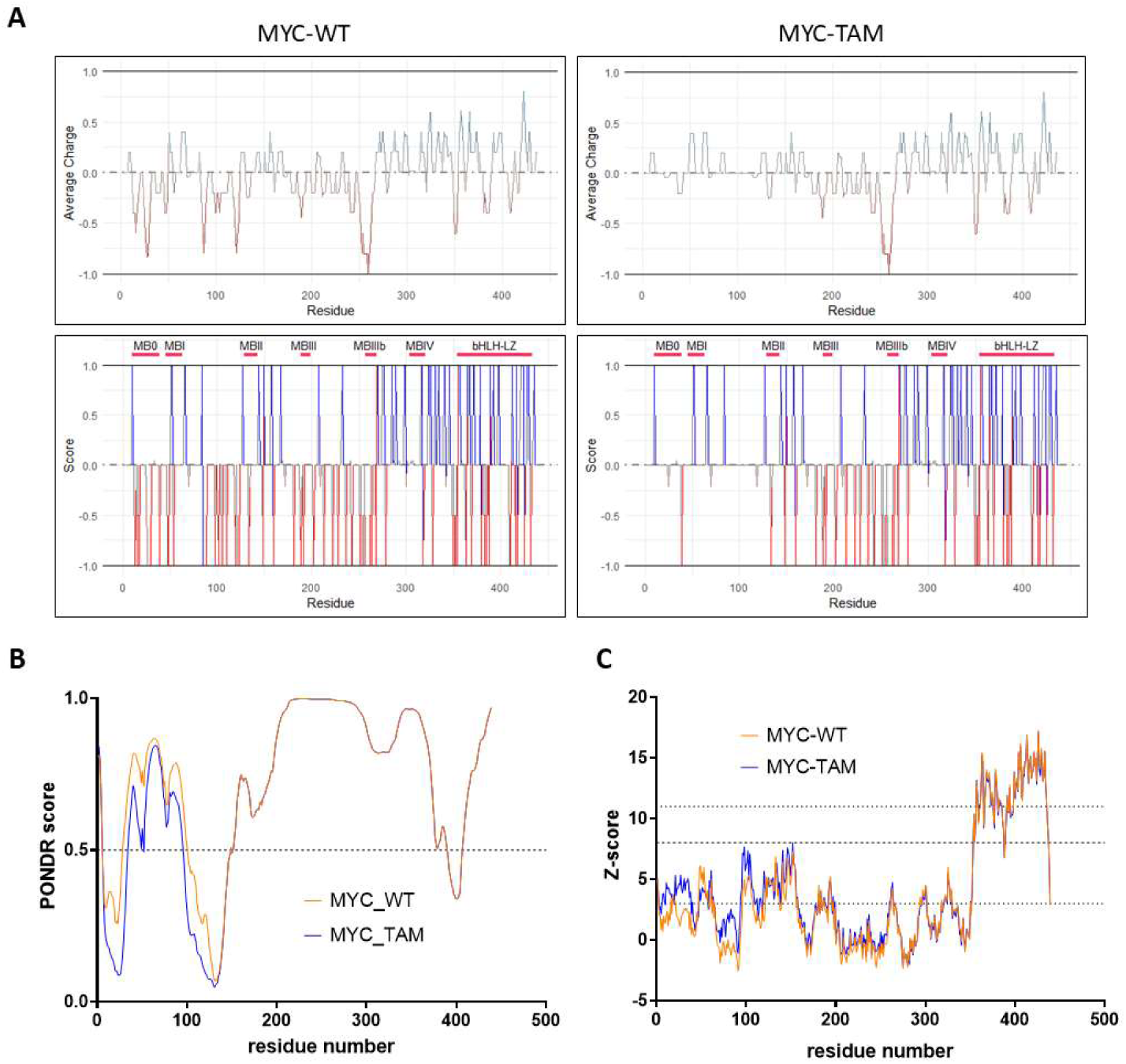
A) Net charge per residue analysis plot for MYC-WT (left) and MYC-TAM (right) using a window size of 5 residues. Positions with an average charge >0 are represented as basic and positions with charge <0 are represented as acidic (top panel). Net charge per residue analysis for MYC-WT (left) and MYC-TAM (right) using a single residue window. Positions with score>0 are predicted as basic whereas position with score<0 are predicted as acidic (low panel). All graphs were generated by R package IDPR. B) Predicted intrinsic disorder for MYC-WT and MYC-TAM using PONDR (Predictor of Natural Disordered Regions) VSL2 score (y axis), and amino acid positions (x axis)(PONDR score>0.5 indicates disorder). C) Predicted intrinsic disorder for MYC-WT and MYC-TAM using ADOPT (82). ADOPT score>8 indicates order, 3<ADOPT score<8 indicates partial disorder, and ADOPT score<3 indicates full disorder.

**Supplementary figure 8:**
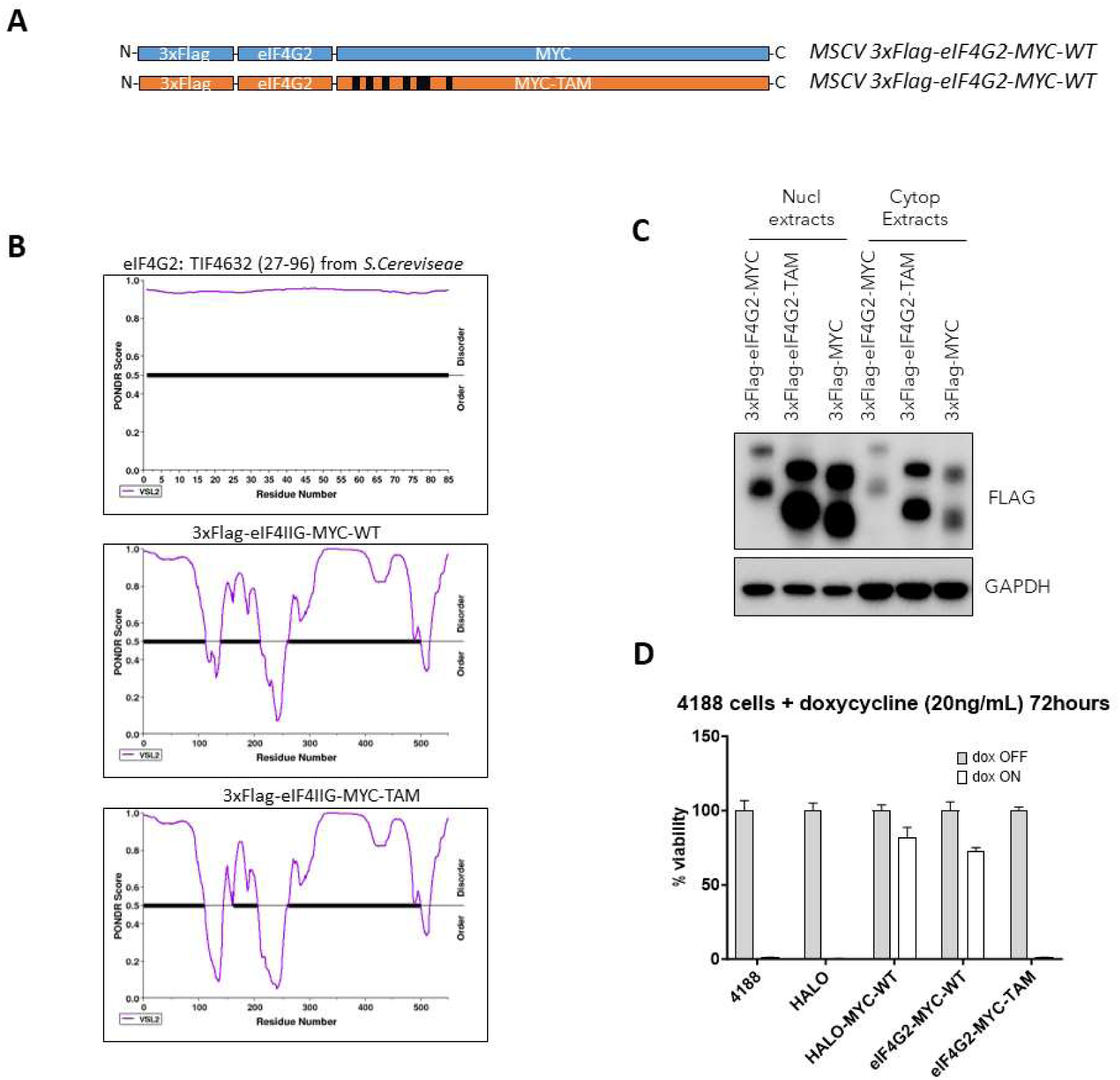
A) Schematic of eIF4G2-tagged constructs tested. The eIF4G2 moiety corresponds to the TIF4632_27-96_ from *S. Cereviseae.* B) Plots depicting intrinsic disorder for eIF4G2 (top), 3xFlag-eIF4G2-MYC-WT (middle) and 3xFlag-eIF4G2-MYC-TAM (bottom). PONDR (Predictor of Natural Disordered Regions) VSL2 score is shown on the y axis, and amino acid positions are shown on the x axis. C) 4188 cells were transduced to stably express 3xFlag-eIF4G2-MYC-WT and 3xFlag-eIF4G2-MYC-TAM. Nuclear and cytoplasmic extracts were immunoblotted for Flag expression. Lysates from cells expressing 3xFlag-MYC were used as positive control. GAPDH was used as loading control. D) Cell viability of 4188 cells after incubation with doxycycline 72h with doxycycline (20ng/mL). Cells were stably transduced to express 3xFlag-eIF4G2-MYC-WT and 3xFlag-eIF4G2-MYC-TAM. Cells expressing HALO and cells expressing HALO-MYC-WT were used as negative and as positive control, respectively.

**Supplementary figure 9:**
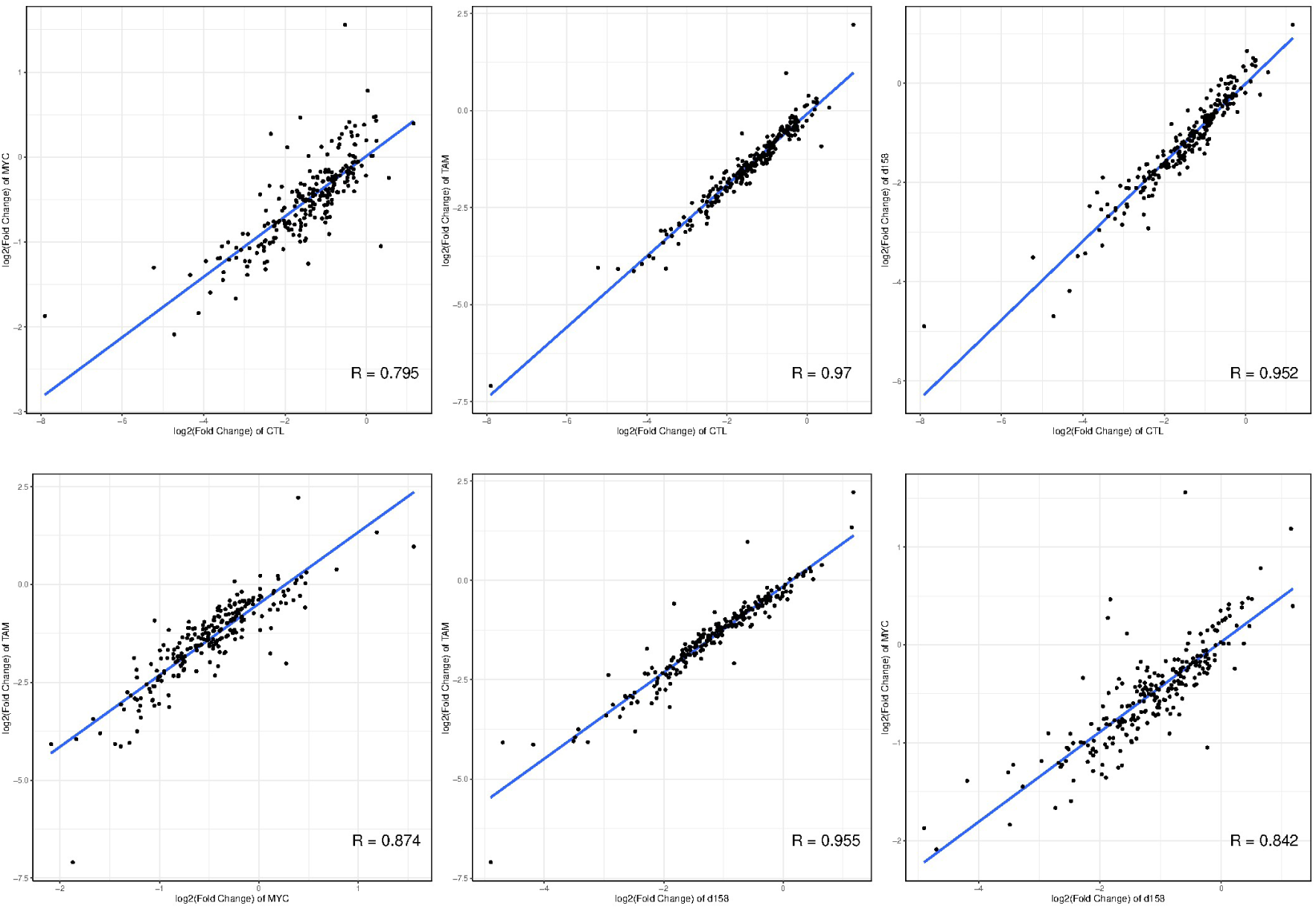
Pearson’s correlation of log2 fold-change between RNA-seq for significantly differentially deregulated genes after doxycycline incubation compared to no-doxycycline condition. CTL: control un-transfected 4188 cells; MYC: 4188 cells stably expressing ectopic MYC-WT; TAM: : 4188 cells stably expressing ectopic MYC-TAM; d158: 4188 cells stably expressing ectopic Δ158-MYC mutant.

**Supplementary figure 10:**
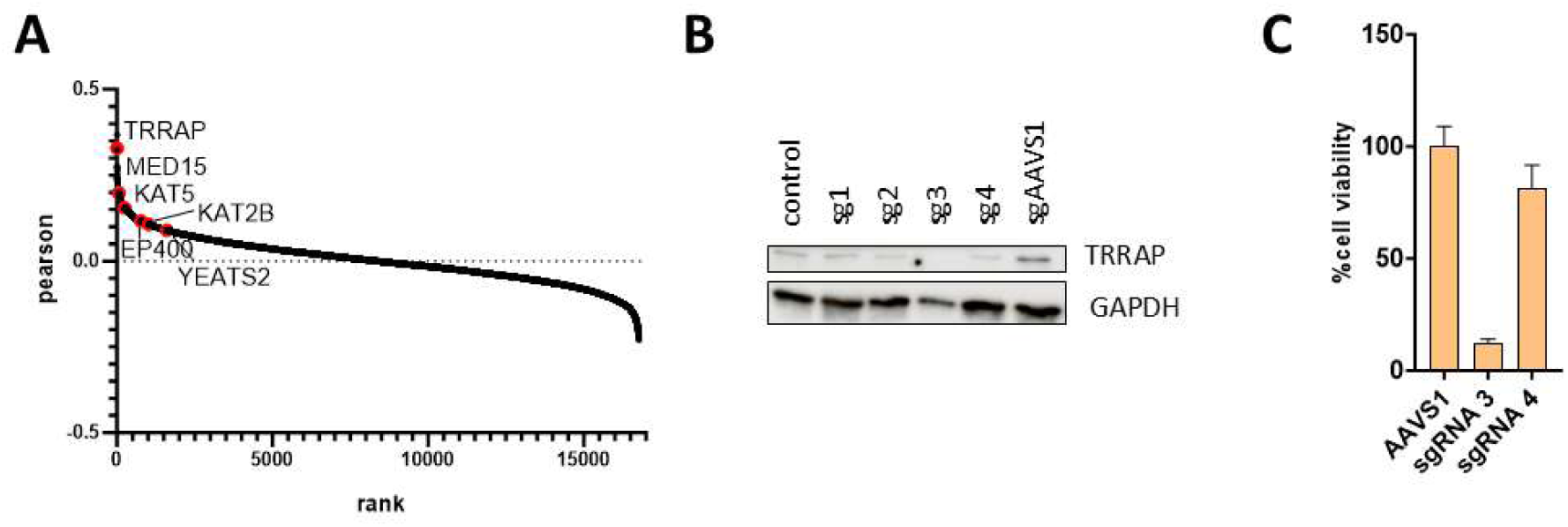
A) Waterfall plot showing the Pearson correlation of proteins from DepMap database with MYC and ranked by Pearson coefficient. Table shows the MYC cofactors identified by BioID MS that show the higher Pearson coefficient. The top 6 identified MYC co-factors are highlighted in red in the waterfall plot. B) Immunoblot of TRRAP expression following transfection with CRISPR guide RNAs (day 7 post-transduction, n=1). AAVS1: guide targeting the safe-harbour locus as control. C) Cell viability measured by Cell-Titter Glo after transfection with guides #3 and #4 (sg3 and sg4, respectively) or with AAVS1 (day 5 post-transduction, n=1).

**Supplementary figure 11:**
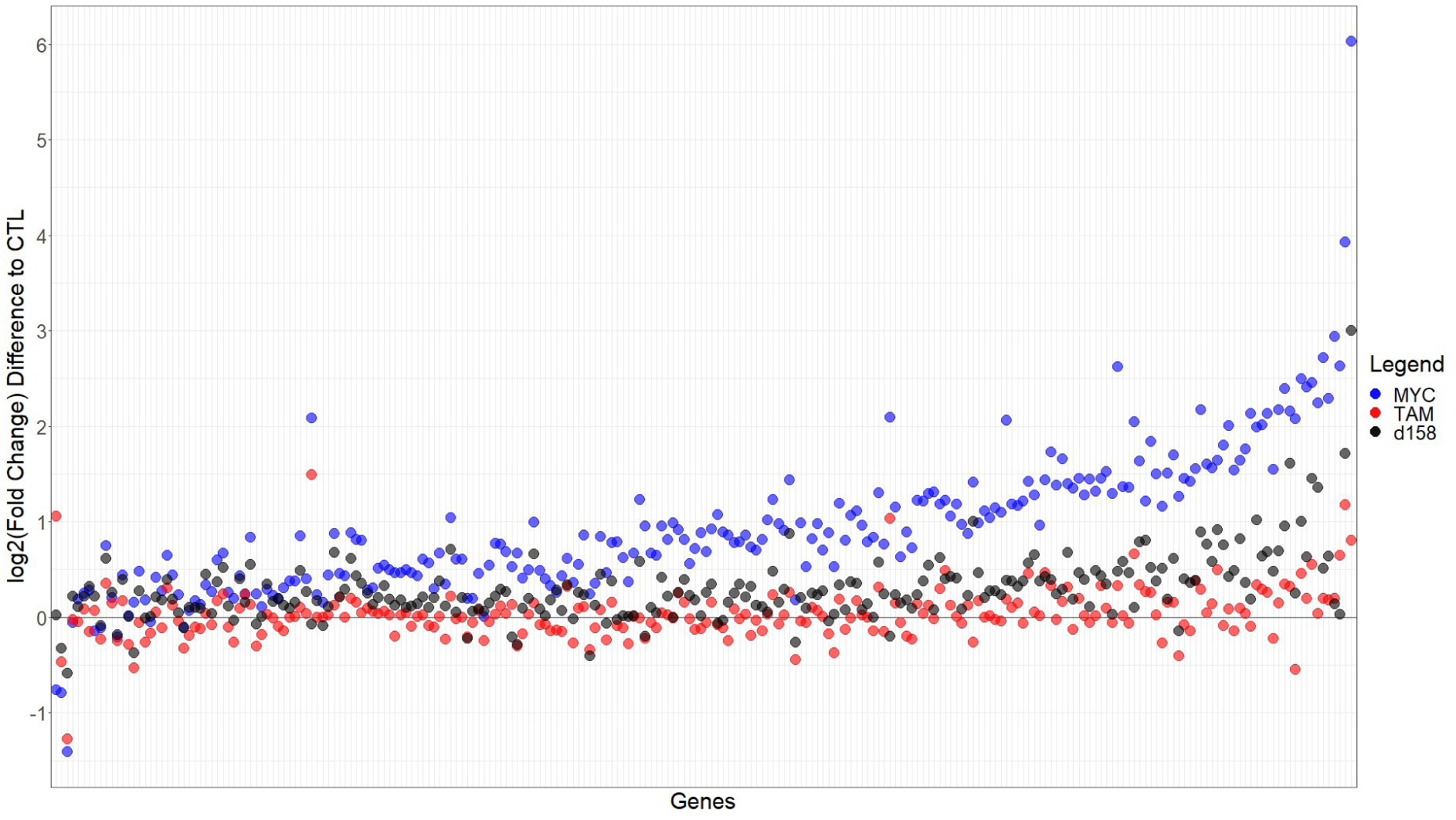
Dot plot showing the changes in gene expression as difference in log2 Fold Change compared to control after doxycycline incubation. Genes are sorted in descending order according to the log2 Fold Change value between dox^on^ vs dox^off^ conditions observed in the control non-transduced cells. For a given gene, the larger the change in expression is in control, the more significant recovery of gene expression is observed in cells expressing ectopic MYC. MYC: 4188 cells stably expressing ectopic MYC-WT; TAM: : 4188 cells stably expressing ectopic MYC-TAM; d158: 4188 cells stably expressing ectopic Δ158-MYC mutant.

## REFERENCES

1. Prendergast GC. Mechanisms of apoptosis by c-Myc. Oncogene. 1999;18(19):2967–87.

2. Dang CV. MYC, metabolism, cell growth, and tumorigenesis. Cold Spring Harb Perspect Med. 2013;3(8).

3. Bretones G, Delgado MD, Leon J. Myc and cell cycle control. Biochim Biophys Acta. 2015;1849(5):506–16.

4. Henriksson M, Luscher B. Proteins of the Myc network: essential regulators of cell growth and differentiation. Adv Cancer Res. 1996;68:109–82.

5. Huber AL, Papp SJ, Chan AB, Henriksson E, Jordan SD, Kriebs A, et al. CRY2 and FBXL3 Cooperatively Degrade c-MYC. Mol Cell. 2016;64(4):774–89.

6. Hanahan D, Weinberg RA. Hallmarks of cancer: the next generation. Cell. 2011;144(5):646–74.

7. Gabay M, Li Y, Felsher DW. MYC activation is a hallmark of cancer initiation and maintenance. Cold Spring Harb Perspect Med. 2014;4(6).

8. Carabet LA, Rennie PS, Cherkasov A. Therapeutic Inhibition of Myc in Cancer. Structural Bases and Computer-Aided Drug Discovery Approaches. Int J Mol Sci. 2018;20(1).

9. Beroukhim R, Mermel CH, Porter D, Wei G, Raychaudhuri S, Donovan J, et al. The landscape of somatic copy-number alteration across human cancers. Nature. 2010;463(7283):899–905.

10. Duffy MJ, O’Grady S, Tang M, Crown J. MYC as a target for cancer treatment. Cancer Treat Rev. 2021;94:102154.

11. Weng AP, Ferrando AA, Lee W, Morris JPt, Silverman LB, Sanchez-Irizarry C, et al. Activating mutations of NOTCH1 in human T cell acute lymphoblastic leukemia. Science. 2004;306(5694):269-71.

12. Herranz D, Ferrando AA. An oncogenic enhancer enemy (N-Me) in T-ALL. Cell Cycle. 2015;14(2):167–8.

13. Sanchez-Martin M, Ferrando A. The NOTCH1-MYC highway toward T-cell acute lymphoblastic leukemia. Blood. 2017;129(9):1124–33.

14. Delgado MD, Leon J. Myc roles in hematopoiesis and leukemia. Genes Cancer. 2010;1(6):605–16.

15. Raetz EA, Teachey DT. T-cell acute lymphoblastic leukemia. Hematology Am Soc Hematol Educ Program. 2016;2016(1):580–8.

16. Soucek L, Whitfield J, Martins CP, Finch AJ, Murphy DJ, Sodir NM, et al. Modelling Myc inhibition as a cancer therapy. Nature. 2008;455(7213):679-83.

17. Soucek L, Whitfield JR, Sodir NM, Masso-Valles D, Serrano E, Karnezis AN, et al. Inhibition of Myc family proteins eradicates KRas-driven lung cancer in mice. Genes Dev. 2013;27(5):504–13.

18. King B, Trimarchi T, Reavie L, Xu L, Mullenders J, Ntziachristos P, et al. The ubiquitin ligase FBXW7 modulates leukemia-initiating cell activity by regulating MYC stability. Cell. 2013;153(7):1552–66.

19. Gartel AL, Ye X, Goufman E, Shianov P, Hay N, Najmabadi F, Tyner AL. Myc represses the p21(WAF1/CIP1) promoter and interacts with Sp1/Sp3. Proc Natl Acad Sci U S A. 2001;98(8):4510–5.

20. Conacci-Sorrell M, McFerrin L, Eisenman RN. An overview of MYC and its interactome. Cold Spring Harb Perspect Med. 2014;4(1):a014357.

21. Thomas LR, Adams CM, Wang J, Weissmiller AM, Creighton J, Lorey SL, et al. Interaction of the oncoprotein transcription factor MYC with its chromatin cofactor WDR5 is essential for tumor maintenance. Proc Natl Acad Sci U S A. 2019;116(50):25260–8.

22. Kenneth NS, Ramsbottom BA, Gomez-Roman N, Marshall L, Cole PA, White RJ. TRRAP and GCN5 are used by c-Myc to activate RNA polymerase III transcription. Proc Natl Acad Sci U S A. 2007;104(38):14917–22.

23. Bayliss R, Burgess SG, Leen E, Richards MW. A moving target: structure and disorder in pursuit of Myc inhibitors. Biochem Soc Trans. 2017;45(3):709–17.

24. Thomas LR, Wang Q, Grieb BC, Phan J, Foshage AM, Sun Q, et al. Interaction with WDR5 promotes target gene recognition and tumorigenesis by MYC. Mol Cell. 2015;58(3):440–52.

25. Richards MW, Burgess SG, Poon E, Carstensen A, Eilers M, Chesler L, Bayliss R. Structural basis of N-Myc binding by Aurora-A and its destabilization by kinase inhibitors. Proc Natl Acad Sci U S A. 2016;113(48):13726–31.

26. Wright PE, Dyson HJ. Intrinsically disordered proteins in cellular signalling and regulation. Nat Rev Mol Cell Biol. 2015;16(1):18–29.

27. Felsher DW, Bishop JM. Reversible tumorigenesis by MYC in hematopoietic lineages. Mol Cell. 1999;4(2):199–207.

28. Tansey. Mammalian MYC Proteins and Cancer. New Journal of Science. 2014;Volume 2014.

29. Kato GJ, Barrett J, Villa-Garcia M, Dang CV. An amino-terminal c-myc domain required for neoplastic transformation activates transcription. Mol Cell Biol. 1990;10(11):5914–20.

30. Rabellino A, Melegari M, Tompkins V, Chen WN, Van Ness BG, Teruya-Feldstein J, et al. PIAS1 Promotes Lymphomagenesis through MYC Upregulation. Cell Reports. 2016;15(10):2266–78.

31. De Melo J, Kim SS, Lourenco C, Penn LZ. Lysine-52 stabilizes the MYC oncoprotein through an SCF(Fbxw7)-independent mechanism. Oncogene. 2017;36(49):6815–22.

32. Smith MJ, Charron-Prochownik DC, Prochownik EV. The leucine zipper of c-Myc is required for full inhibition of erythroleukemia differentiation. Mol Cell Biol. 1990;10(10):5333–9.

33. Frietze S, Farnham PJ. Transcription factor effector domains. Subcell Biochem. 2011;52:261–77.

34. Theillet FX, Kalmar L, Tompa P, Han KH, Selenko P, Dunker AK, et al. The alphabet of intrinsic disorder: I. Act like a Pro: On the abundance and roles of proline residues in intrinsically disordered proteins. Intrinsically Disord Proteins. 2013;1(1):e24360.

35. Uversky VN. The intrinsic disorder alphabet. III. Dual personality of serine. Intrinsically Disord Proteins. 2015;3(1):e1027032.

36. Sanborn AL, Yeh BT, Feigerle JT, Hao CV, Townshend RJ, Lieberman Aiden E, et al. Simple biochemical features underlie transcriptional activation domain diversity and dynamic, fuzzy binding to Mediator. Elife. 2021;10.

37. Erijman A, Kozlowski L, Sohrabi-Jahromi S, Fishburn J, Warfield L, Schreiber J, et al. A High-Throughput Screen for Transcription Activation Domains Reveals Their Sequence Features and Permits Prediction by Deep Learning. Mol Cell. 2020;79(6):1066.

38. Yang J, Zeng Y, Liu Y, Gao M, Liu S, Su Z, Huang Y. Electrostatic interactions in molecular recognition of intrinsically disordered proteins. J Biomol Struct Dyn. 2020;38(16):4883–94.

39. Shi B, Li W, Song Y, Wang Z, Ju R, Ulman A, et al. UTX condensation underlies its tumour-suppressive activity. Nature. 2021;597(7878):726-31.

40. Langenau DM, Traver D, Ferrando AA, Kutok JL, Aster JC, Kanki JP, et al. Myc-induced T cell leukemia in transgenic zebrafish. Science. 2003;299(5608):887–90.

41. Mansour MR, He S, Li Z, Lobbardi R, Abraham BJ, Hug C, et al. JDP2: An oncogenic bZIP transcription factor in T cell acute lymphoblastic leukemia. J Exp Med. 2018;215(7):1929–45.

42. Zeller KI, Jegga AG, Aronow BJ, O’Donnell KA, Dang CV. An integrated database of genes responsive to the Myc oncogenic transcription factor: identification of direct genomic targets. Genome Biol. 2003;4(10):R69.

43. Farrell AS, Sears RC. MYC degradation. Cold Spring Harb Perspect Med. 2014;4(3).

44. Small GW, Chou TY, Dang CV, Orlowski RZ. Evidence for involvement of calpain in c-Myc proteolysis in vivo. Arch Biochem Biophys. 2002;400(2):151–61.

45. Llombart V, Mansour MR. Therapeutic targeting of “undruggable” MYC. EBioMedicine. 2022;75:103756.

46. Hinze L, Pfirrmann M, Karim S, Degar J, McGuckin C, Vinjamur D, et al. Synthetic Lethality of Wnt Pathway Activation and Asparaginase in Drug-Resistant Acute Leukemias. Cancer Cell. 2019;35(4):664–76 e7.

47. Brown CE, Howe L, Sousa K, Alley SC, Carrozza MJ, Tan S, Workman JL. Recruitment of HAT complexes by direct activator interactions with the ATM-related Tra1 subunit. Science. 2001;292(5525):2333-7.

48. Grant PA, Schieltz D, Pray-Grant MG, Yates JR, 3rd, Workman JL. The ATM-related cofactor Tra1 is a component of the purified SAGA complex. Mol Cell. 1998;2(6):863–7.

49. Nikiforov MA, Chandriani S, Park J, Kotenko I, Matheos D, Johnsson A, et al. TRRAP-dependent and TRRAP-independent transcriptional activation by Myc family oncoproteins. Mol Cell Biol. 2002;22(14):5054–63.

50. Feris EJ, Hinds JW, Cole MD. Formation of a structurally-stable conformation by the intrinsically disordered MYC:TRRAP complex. PLoS One. 2019;14(12):e0225784.

51. Liu X, Tesfai J, Evrard YA, Dent SY, Martinez E. c-Myc transformation domain recruits the human STAGA complex and requires TRRAP and GCN5 acetylase activity for transcription activation. J Biol Chem. 2003;278(22):20405–12.

52. Zhang N, Ichikawa W, Faiola F, Lo SY, Liu X, Martinez E. MYC interacts with the human STAGA coactivator complex via multivalent contacts with the GCN5 and TRRAP subunits. Biochim Biophys Acta. 2014;1839(5):395–405.

53. Park J, Kunjibettu S, McMahon SB, Cole MD. The ATM-related domain of TRRAP is required for histone acetyltransferase recruitment and Myc-dependent oncogenesis. Gene Dev. 2001;15(13):1619–24.

54. Janicki SM, Tsukamoto T, Salghetti SE, Tansey WP, Sachidanandam R, Prasanth KV, et al. From silencing to gene expression: real-time analysis in single cells. Cell. 2004;116(5):683–98.

55. Helander S, Montecchio M, Pilstal R, Su Y, Kuruvilla J, Elven M, et al. Pre-Anchoring of Pin1 to Unphosphorylated c-Myc in a Fuzzy Complex Regulates c-Myc Activity. Structure. 2015;23(12):2267–79.

56. Amin C, Wagner AJ, Hay N. Sequence-specific transcriptional activation by Myc and repression by Max. Mol Cell Biol. 1993;13(1):383–90.

57. Stone J, de Lange T, Ramsay G, Jakobovits E, Bishop JM, Varmus H, Lee W. Definition of regions in human c-myc that are involved in transformation and nuclear localization. Mol Cell Biol. 1987;7(5):1697–709.

58. Kalkat M, Resetca D, Lourenco C, Chan PK, Wei Y, Shiah YJ, et al. MYC Protein Interactome Profiling Reveals Functionally Distinct Regions that Cooperate to Drive Tumorigenesis. Mol Cell. 2018;72(5):836–48 e7.

59. Cowling VH, Cole MD. Mechanism of transcriptional activation by the Myc oncoproteins. Semin Cancer Biol. 2006;16(4):242–52.

60. Zhao LJ, Loewenstein PM, Green M. Enhanced MYC association with the NuA4 histone acetyltransferase complex mediated by the adenovirus E1A N-terminal domain activates a subset of MYC target genes highly expressed in cancer cells. Genes Cancer. 2017;8(11-12):752–61.

61. Vassilev A, Yamauchi J, Kotani T, Prives C, Avantaggiati ML, Qin J, Nakatani Y. The 400 kDa subunit of the PCAF histone acetylase complex belongs to the ATM superfamily. Mol Cell. 1998;2(6):869–75.

62. Poole CJ, van Riggelen J. MYC-Master Regulator of the Cancer Epigenome and Transcriptome. Genes (Basel). 2017;8(5).

63. McMahon SB, Van Buskirk HA, Dugan KA, Copeland TD, Cole MD. The novel ATM-related protein TRRAP is an essential cofactor for the c-Myc and E2F oncoproteins. Cell. 1998;94(3):363–74.

64. Sigler PB. Transcriptional activation. Acid blobs and negative noodles. Nature. 1988;333(6170):210-2.

65. Tuttle LM, Pacheco D, Warfield L, Luo J, Ranish J, Hahn S, Klevit RE. Gcn4-Mediator Specificity Is Mediated by a Large and Dynamic Fuzzy Protein-Protein Complex. Cell Reports. 2018;22(12):3251–64.

66. Sabari BR, Dall’Agnese A, Boija A, Klein IA, Coffey EL, Shrinivas K, et al. Coactivator condensation at super-enhancers links phase separation and gene control. Science. 2018;361(6400).

67. Boija A, Klein IA, Sabari BR, Dall’Agnese A, Coffey EL, Zamudio AV, et al. Transcription Factors Activate Genes through the Phase-Separation Capacity of Their Activation Domains. Cell. 2018;175(7):1842–55 e16.

68. Joung J, Konermann S, Gootenberg JS, Abudayyeh OO, Platt RJ, Brigham MD, et al. Genome-scale CRISPR-Cas9 knockout and transcriptional activation screening. Nat Protoc. 2017;12(4):828–63.

69. Poliseno L, Mariani L, Collecchi P, Piras A, Zaccaro L, Rainaldi G. Bcl2-negative MCF7 cells overexpress p53: implications for the cell cycle and sensitivity to cytotoxic drugs. Cancer Chemother Pharmacol. 2002;50(2):127–30.

70. Wasylishen AR, Stojanova A, Oliveri S, Rust AC, Schimmer AD, Penn LZ. New model systems provide insights into Myc-induced transformation. Oncogene. 2011;30(34):3727–34.

71. Sanda T, Lawton LN, Barrasa MI, Fan ZP, Kohlhammer H, Gutierrez A, et al. Core transcriptional regulatory circuit controlled by the TAL1 complex in human T cell acute lymphoblastic leukemia. Cancer Cell. 2012;22(2):209–21.

72. Wilkins MR, Gasteiger E, Bairoch A, Sanchez JC, Williams KL, Appel RD, Hochstrasser DF. Protein identification and analysis tools in the ExPASy server. Methods Mol Biol. 1999;112:531–52.

73. Ginell GM, Holehouse AS. Analyzing the Sequences of Intrinsically Disordered Regions with CIDER and localCIDER. Methods Mol Biol. 2020;2141:103–26.

74. Barshop WD, Kim HJ, Fan X, Sha J, Rayatpisheh S, Wohlschlegel JA. Chemical Derivatization of Affinity Matrices Provides Protection from Tryptic Proteolysis. J Proteome Res. 2019;18(10):3586–96.

75. Cox J, Mann M. MaxQuant enables high peptide identification rates, individualized p.p.b.-range mass accuracies and proteome-wide protein quantification. Nat Biotechnol. 2008;26(12):1367–72.

76. Choi M, Chang CY, Clough T, Broudy D, Killeen T, MacLean B, Vitek O. MSstats: an R package for statistical analysis of quantitative mass spectrometry-based proteomic experiments. Bioinformatics. 2014;30(17):2524–6.

77. Benjamini Y, Hochberg Y. Controlling the False Discovery Rate - a Practical and Powerful Approach to Multiple Testing. J R Stat Soc B. 1995;57(1):289–300.

78. Szklarczyk D, Franceschini A, Wyder S, Forslund K, Heller D, Huerta-Cepas J, et al. STRING v10: protein-protein interaction networks, integrated over the tree of life. Nucleic Acids Res. 2015;43(Database issue):D447–52.

79. Kappei D, Scheibe M, Paszkowski-Rogacz M, Bluhm A, Gossmann TI, Dietz S, et al. Phylointeractomics reconstructs functional evolution of protein binding. Nat Commun. 2017;8:14334.

80. Pertea M, Pertea GM, Antonescu CM, Chang TC, Mendell JT, Salzberg SL. StringTie enables improved reconstruction of a transcriptome from RNA-seq reads. Nat Biotechnol. 2015;33(3):290–5.

81. Sanson KR, Hanna RE, Hegde M, Donovan KF, Strand C, Sullender ME, et al. Optimized libraries for CRISPR-Cas9 genetic screens with multiple modalities. Nat Commun. 2018;9(1):5416.

82. Redl I, Fisicaro C, Dutton O, Hoffmann F, Henderson L, Owens BMJ, et al. ADOPT: intrinsic protein disorder prediction through deep bidirectional transformers. NAR Genom Bioinform. 2023;5(2):lqad041.

83. Perez-Riverol Y, Bai J, Bandla C, Garcia-Seisdedos D, Hewapathirana S, Kamatchinathan S, et al. The PRIDE database resources in 2022: a hub for mass spectrometry-based proteomics evidences. Nucleic Acids Res. 2022;50(D1):D543–D52.

84. Edgar R, Domrachev M, Lash AE. Gene Expression Omnibus: NCBI gene expression and hybridization array data repository. Nucleic Acids Res. 2002;30(1):207–10.

85. Xie Y, Li H, Luo X, Li H, Gao Q, Zhang L, et al. IBS 2.0: an upgraded illustrator for the visualization of biological sequences. Nucleic Acids Res. 2022;50(W1):W420–W6.

